# Homochiral and racemic MicroED structures of a peptide from the ice nucleation protein InaZ

**DOI:** 10.1101/473116

**Authors:** Chih-Te Zee, Calina Glynn, Marcus Gallagher-Jones, Jennifer Miao, Carlos G. Santiago, Duilio Cascio, Tamir Gonen, Michael R. Sawaya, Jose A. Rodriguez

## Abstract

The ice nucleation protein InaZ of *Pseudomonas syringae* contains a large number of degenerate repeats that span more than a quarter of its sequence and include the segment GSTSTA. We determine *ab initio* structures of this repeat segment, resolved to 1.1Å by microfocus x-ray crystallography and 0.9Å by the cryoEM method MicroED, from both racemic and homochiral crystals. We evaluate the benefits of racemic protein crystals for structure determination by MicroED and confirm that phase restriction introduced by crystal centrosymmetry increases the number of successful trials during *ab initio* phasing of electron diffraction data. Both homochiral and racemic GSTSTA form amyloid-like protofibrils with labile, corrugated antiparallel beta sheets that mate face to back. The racemic GSTSTA protofibril represents a new class of amyloid assembly in which all left-handed sheets mate with their all right-handed counterparts. Our determination of racemic amyloid assemblies by MicroED reveals complex amyloid architectures and illustrates the racemic advantage in macromolecular crystallography, now with sub-micron sized crystals.

**Synopsis:** The atomic asymmetry, left or right handedness, present in macromolecules and first described by Pasteur in his experiments with tartaric acid, is evident even in complex molecular assemblies like amyloid fibrils. Here, using the cryoEM method MicroED, we show that a segment from the ice nucleation protein InaZ assembles into homochiral and racemic water-binding amyloid protofibrils.

## 1. Introduction

Expressed by a subset of microorganisms, ice nucleation proteins are capable of stimulating ice formation in supercooled water (Green & Warren, 1985). The gram-negative microbe, *P. syringae* is sold commercially as Snowmax^®^ for its ice nucleating activity (Green & Warren, 1985; Cochet & Widehem, 2000). The ice nucleation protein InaZ is produced by *Pseudomonas syringae* and localized to its outer membrane (Green & Warren, 1985; Wolber *et al*., 1986). The sequence of InaZ is approximately 1200 residues in length, over half of which includes degenerate octapeptide repeats. A subpopulation of degenerate repeats share the consensus motif GSTXT[A/S], where X represents an unconserved amino acid (Fig. S1) (Green & Warren, 1985; Warren *et al*., 1986). These repeats are shared by other Ina proteins and may collectively contribute to ice nucleation (Green & Warren, 1985; Kobashigawa Yoshihiro *et al*., 2005; Han *et al*., 2017).

Despite the crystallographic determination of structures for other ice binding proteins (Davies, 2014; Garnham, Campbell, & Davies, 2011), InaZ remains recalcitrant to crystallization. Models of full-length InaZ propose it has a beta helical (Garnham, Campbell, Walker *et al*., 2011; Graether & Jia, 2001) or solenoid-like fold, rich in stacked beta strands (Cochet & Widehem, 2000; Pandey *et al*., 2016). These features are shared by amyloid filaments: their tightly mated beta sheets form fibrils that can crosslink, cluster and be functional (Nelson *et al*., 2005; Sawaya *et al*., 2007; Fitzpatrick *et al*., 2017; Eisenberg & Jucker, 2012; Maury, 2009). Functional amyloid assemblies appear across the tree of life (Wasmer *et al*., 2008; Hughes *et al*., 2018; Maury, 2009; Tayeb-Fligelman *et al*., 2017) and can contain low complexity regions with degenerate repeats (Hughes *et al*., 2018).

Success in determining amyloid structures was first achieved by crystallizing short segments that stabilize the cores of fibrils through a motif known as the steric zipper (Nelson *et al*., 2005; Sawaya *et al*., 2007). However, the propensity of elongated beta strands to twist or kink can limit crystal growth, sometimes yielding nanocrystals that pose a challenge for structure determination (Rodriguez *et al*., 2015). These limits have recently been overcome in part by the development of the cryo electron microscopy (cryoEM) method, electron micro diffraction (MicroED) (Shi *et al*., 2013). MicroED yields high-resolution structures from protein crystals no thicker than a few hundred nanometers (Shi *et al*., 2016; Rodriguez *et al*., 2017). Because of this, MicroED has helped determine the structures of a number of amyloid protofibrils (Rodriguez *et al*., 2015; Krotee *et al*.) with atomic resolution; some *ab initio* (Sawaya *et al*., 2016; Gallagher-Jones *et al*., 2018).

Racemic crystallography further facilitates crystallization of proteins and peptides (Matthews, 2009; Yeates & Kent, 2012; Patterson *et al*., 1999), including ice binding proteins (Pentelute *et al*., 2008). Mixing left (L) and right (D) handed enantiomers of a macromolecule improves its likelihood of crystallization and facilitates structural analysis (Yeates & Kent, 2012; Wukovitz & Yeates, 1995). Crystallographic phases are restricted for data from centrosymmetric crystals, making the phase problem associated with the determination of their structure more tractable (Yeates & Kent, 2012). This is advantageous for structure determination by direct methods (Hauptman, 1986), where phases must be computed from measured intensities alone (Hauptman, 1986, 2001; Sheldrick, 2008). Accordingly, various polypeptide structures have been determined by racemic x-ray crystallography, including those of ester insulin, plectasin, and an antifreeze protein (Pentelute *et al*., 2008; Avital-Shmilovici *et al*., 2013; Mandal *et al*., 2009, 2012). While the benefits of racemic crystallography are evident for x-ray diffraction (Matthews, 2009), questions remain about the potential for exploiting these benefits by MicroED.

Hypothesizing that repeat segments of the ice nucleation protein InaZ may form amyloid-like assemblies, we set out to interrogate the structure of GSTSTA from both homochiral and racemic crystals by MicroED. In doing so, we also assessed the fidelity of MicroED data in racemic structure determination. Comparing the structures of homochiral and racemic GSTSTA, we gauge the effect of racemic self-assembly on protofibril architecture. With these structures of a core repeat in the InaZ protein, we begin an atomic level investigation of peptide segments derived from ice nucleation proteins (Pandey *et al*., 2016).

## 2. Results

### 2.1. Identification, synthesis and crystallization of amyloid forming INP segments

With the goal of characterizing the structural properties of degenerate repeats in INPs, we identified a group of hexapeptides within the set of InaZ repeats and evaluated their amyloid forming propensity (Goldschmidt *et al*., 2010) (Fig. S1). We ranked the hexapeptides based on their predicted propensity for amyloid zipper formation, their repeated appearance in INP sequences, and whether they contained polar residues including threonine (Fig. S1). We chose to further investigate a segment whose sequence, GSTSTA, appears identically five times within InaZ: residues 707-712, 755-760, 803-808, 851-856, and 899-904. For simplicity, we numbered the segment 707-712.

We evaluated the crystallization potential of synthesized L- and D-enantiomers of GSTSTA (Fig. S2) compared to that of their racemic mixture by performing high-throughput crystallization trials and monitoring crystal growth. Most crystals appeared within two weeks of the start of each trial. In some conditions, crystallization was observed as early as three hours after the start of the trial. Racemic mixtures produced a greater number of successful crystallization conditions across a broad variety of trials (Fig. S3). The number of successful conditions identified for racemic mixtures outpaced those identified for each enantiomer alone (Fig. S3), consistent with previous predictions (Yeates & Kent, 2012). In conditions where both racemic and single enantiomer crystals grew, racemic crystals appeared sooner (Fig. S3). Minor differences in the speed of crystal appearance and the total number of conditions with identifiable crystals were also seen between L- and D-enantiomers. Fewer conditions were found to grow D-enantiomer crystals across all trials (Fig. S3). These differences may have been a consequence of subtle inequities in the amount of residual trifluoroacetic acid (TFA) associated with each enantiomer in lyophilized powders. Those effects may have been magnified by the relatively high concentrations of peptide required for crystallization of these segments (~100 to 150 mM).

The crystallization conditions chosen for structure determination of homochiral (L-) and racemic (DL-) GSTSTA yielded a high density of well-ordered microcrystals, each with a unique powder diffraction pattern indicating they had formed distinct structures (Fig. S4). Microcrystals were optimized from these conditions for microfocal x-ray diffraction; unoptimized batch conditions yielded nanocrystal slurries directly suitable for MicroED. Since powder diffraction patterns of homochiral GSTSTA crystals were identical for both enantiomers (Fig. S4), we focused our investigation on the L-enantiomer.

### 2.2. *Ab initio* structure determination of L-GSTSTA

We optimized crystals of L-GSTSTA for microfocal x-ray diffraction starting from dense needle clusters and ending with single needles (Fig. S5). Crystals grown in batch were monodisperse rods 1-10 micrometers long and 100 to 500 nanometers wide; these diffracted to approximately 0.9 Å by MicroED (Fig. 1). X-ray diffraction from a single crystal of L-GSTSTA yielded a 91.7% complete dataset to approximately 1.1Å resolution (Table S1), while datasets from three crystals of L-GSTSTA obtained by MicroED were merged to achieve a dataset with an overall completeness of 86.4% at 0.9Å resolution. It is important to note that the x-ray data in this case was limited by detector geometry, which could be adjusted to facilitate slightly higher resolution. Atomic structure solutions were determined for L-GSTSTA from both microfocal x-ray and MicroED data by direct methods (Sheldrick, 2008) (Fig. S6).

**Figure 1.**
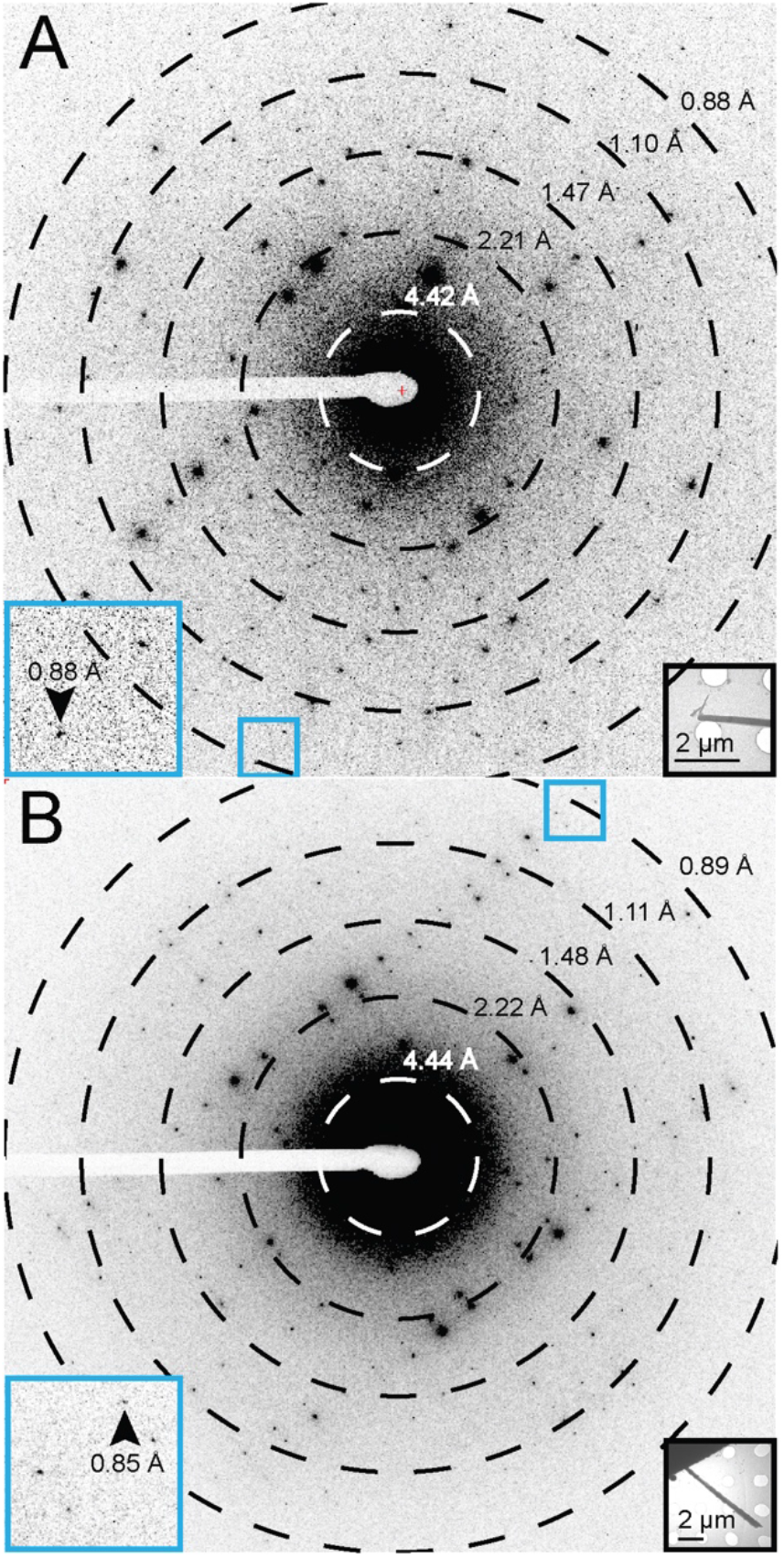
Single diffraction patterns of homochiral L-GSTSTA (A) or racemic GSTSTA (B) measured during continuous rotation MicroED data collection. Each pattern corresponds to a 0.6° wedge (A) or 0.9° wedge (B) of reciprocal space. Black insets show overfocused diffraction images of the crystals used for diffraction; blue squares correspond to magnified regions (blue insets) of the pattern that show diffraction at sub 0.9Å resolution (black arrows). Resolution circles are indicated by rings; scale bars 2μm.

After 50,000 trials, SHELXD yielded correlation figures of merit (CFOM) greater than 80 for both x-ray diffraction and MicroED data (Fig. S6) (Sawaya *et al*., 2016). The initial L-GSTSTA solution with the highest CFOM shows 33 atoms for the x-ray dataset and 36 atoms for the MicroED dataset (Fig. 2A and Fig. S7). During refinement, the number of atoms in the x-ray structure increased to 36 peptide atoms and one bound water (Fig. S7). The final solution achieved from 0.9Å resolution MicroED data also contained 36 atoms in the peptide chain and one water molecule (Fig. 2A).

**Figure 2.**
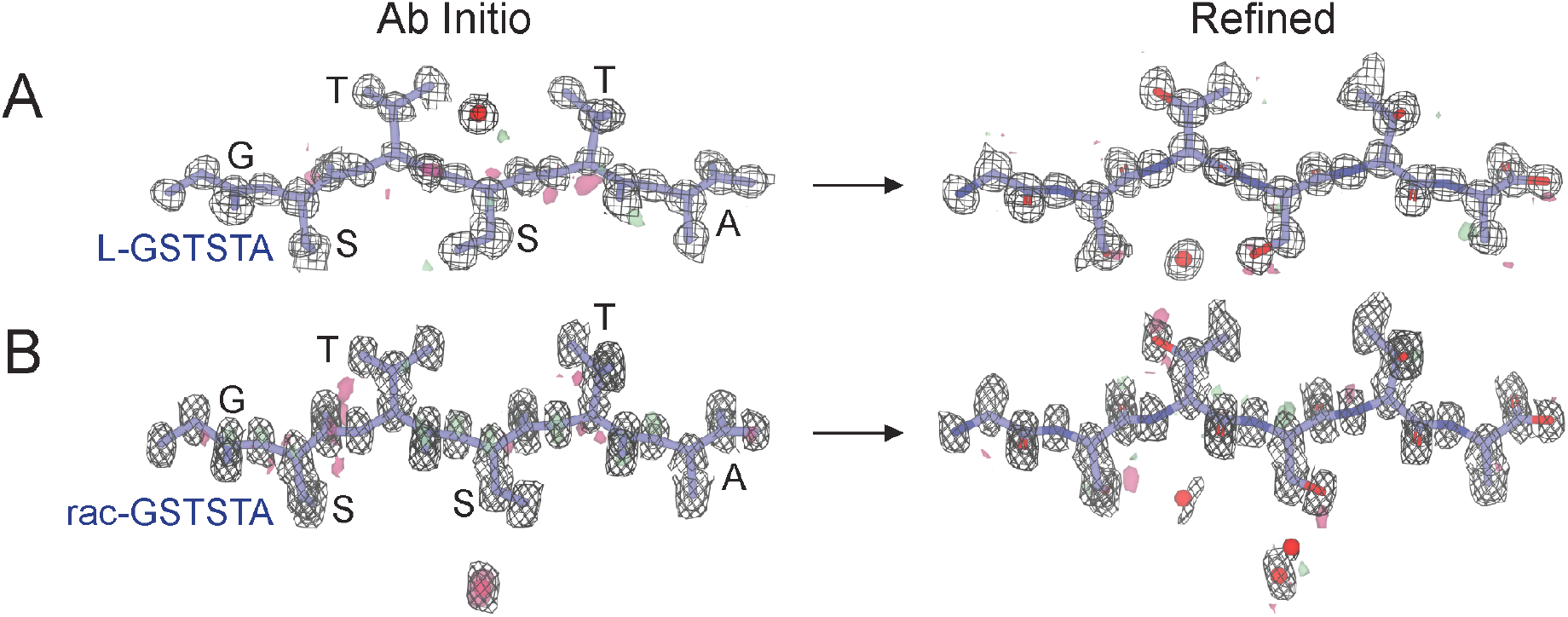
*Ab Initio* structures and electrostatic potential maps of L-GSTSTA (A) or racemic GSTSTA (B). Each map is in A is overlaid onto the initial atomic coordinates calculated by SHELXD from MicroED data. Each map in B is overlaid onto its corresponding refined model. The 2F_o_-F_c_ map represented by the black mesh is contoured at 1.2 σ. Green and red surfaces represent the F_o_-F_c_ maps contoured at 3.0 and −3.0 σ. Modelled waters are present as red spheres. The waters modelled in the *ab initio* solution in A and the refined structure in B are related by symmetry.

### 2.3. *Ab initio* structure determination of racemic GSTSTA from centrosymmetric crystals

Like the enantiomerically pure crystals of GSTSTA, crystals of racemic GSTSTA started as dense needle clusters and were optimized to single needles and diffracted as single crystals on a microfocal x-ray source (Fig. S5). Batch crystals of racemic GSTSTA were also rod shaped, several microns long and a few hundred nanometers in thickness (Fig. 1). These were immediately suitable for MicroED and diffracted to approximately 0.9 Å (Fig. 1). Data from a single crystal obtained by x-ray diffraction produced a 93.7% complete dataset at 1.1Å resolution, while MicroED data from two nanocrystals of racemic GSTSTA were merged to reach an overall completeness of 77.4% at 0.9Å resolution (Table S1). Initial atomic structure solutions for racemic GSTSTA were obtained by direct methods (Sheldrick, 2008) (Fig. 2B and Fig. S7).

As with L-GSTSTA, solutions for the racemic crystals yielded correlation figures of merit (CFOM) greater than 80 after 50,000 trials (Fig. S6). A comparison of racemic GSTSTA to L-GSTSTA datasets indicated a higher number of potentially correct solutions found with racemic GSTSTA data (Fig. S6). MicroED data shows a distribution of CFOM values that is shifted toward higher values, even when truncated to 1.1Å resolution to match the resolution of the x-ray datasets. However, the most dramatic shift in this distribution is evident at 0.9Å resolution (Fig. S6).

Initial solutions with the highest CFOM show a total of 35 peptide atoms and 4 waters, for the structure determined from x-ray data, and a total of 36 peptide atoms and 1 water for that determined by MicroED (Fig. 2B). During refinement, the number of peptide atoms in the x-ray structure increased to 36 (Fig. S7), while the MicroED structure gained two waters (Fig. 2B). Linear regression of observed to calculated structure factors in MicroED data shows an R-value of 0.94 and a slope of 0.97 for data reduced in space group P1 (Fig. 3C). These numbers are in good agreement with those obtained by microfocal x-ray diffraction (Fig. 11C) and indicate a good fit between model and measurement for the racemic GSTSTA structure.

**Figure 3.**
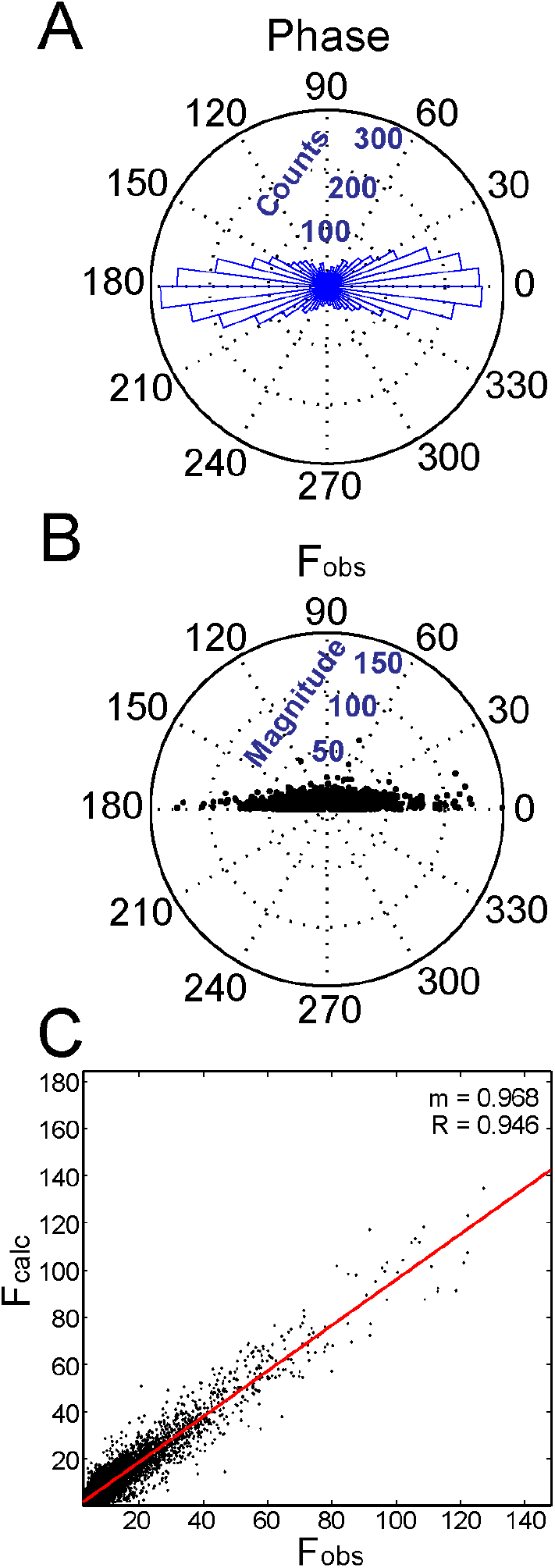
(A) The calculated phase associated with each reflection in the P1 refinement of racemic GSTSTA data obtained by MicroED was analyzed and plotted as a histogram along the unit circle. (B) The magnitude of each reflection is plotted as a function of the absolute value of its associated phase. (C) A plot of F_o_ vs. F_c_ values for each reflection in this data set shows a distribution that can be fit by linear regression, shown as a red line with slope m=0.97 and R-value 0.95.

### 2.4. Paired comparison of Fourier magnitudes measured by x-ray crystallography or MicroED

A comparison between x-ray and MicroED datasets for homochiral crystals of GSTSTA shows these two types of measurement are in close agreement (Fig. S8, S9), albeit slightly higher merge errors are observed in MicroED data across resolution bins (Table S2). A direct comparison of Fourier magnitudes for paired reflections between these datasets is fit by a line with slope m=0.921 and an R-value of 0.826 (Fig. S8). In contrast, the comparison between x-ray and MicroED data for racemic GSTSTA data shows a greater difference between the two sets and a lower R-value for the best fit line comparing Fourier magnitudes of paired x-ray and MicroED reflections (Fig. S8, S10). This difference is likely due to a lack of isomorphism between the cells of racemic GSTSTA crystals used for MicroED data collection vs. x-ray data collection. The cell constants for racemic GSTSTA crystals diffracted by MicroED and microfocal x-ray crystallography were {15.23 9.29 21.06, 90.0 108.2 90.0} and {14.03 9.22 20.77, 90.0 104.5 90.0} respectively (Table S1).

### 2.5. Phase restriction in centrosymmetric crystals evaluated by MicroED

Data from racemic GSTSTA crystals obtained by MicroED and reduced in the centrosymmetric space group P21/c satisfies refinement with imposed phases of 0 or 180 degrees. Refinement of data from the same crystals reduced in space group P1 results in similar residuals to those obtained for space group P21/c (Table S1). Phases that result from refinement of this structure against data reduced in space group P1 appear bilaterally distributed around 0 and 180 degrees (Fig. 3A). Collapse of this bimodal phase distribution around nπ yields a standard deviation of 34.3 degrees (Fig. 3). When the same procedure is applied to data collected from racemic GSTSTA crystals by x-ray diffraction, a similar trend appears: a normal distribution around nπ with a standard deviation of 34.4 degrees (Fig. S11). Bragg reflections that appear in disallowed regions of phase space (90 and 270 degrees) for both MicroED and x-ray diffraction data are generally weakest (Fig. 3 and S11). This suggests that the primary source of phase error in MicroED data, as with x-ray diffraction, may come from noisy or weak reflections.

### 2.6. Structure of L-GSTSTA

L-GSTSTA assembles into anti-parallel in register beta sheets that mate to form a protofibril (Fig. 4A. S12, S13). Sheets are buckled, compressing the fibril along its length with strands spaced approximately 4.6Å apart (Fig. 4A, S14), closer than the typical 4.7–4.8Å spacings seen in amyloid protofibrils (Sawaya *et al*., 2007). This spacing equates to half the L-GSTSTA cell edge along the a-axis, approximately 9.2Å (Table S1). To accommodate this compression, strands tilt approximately 17 degrees with respect to the fibril axis in alternating directions along a sheet, allowing amides to lie askew from the fibril axis (Fig. S14) while maintaining hydrogen bonding along the protofibril axis (Table S3). Sidechains between neighboring sheets tightly interdigitate to create a close packing within the fibril (Fig. S12); inter-sheet distances range from 5 to 7 Å. The interface created at the fibril core is small with 229 Å^2^ of buried surface area but shows a relatively high degree of shape complementarity (S_c_=0.75) (Lawrence & Colman, 1993). The L-GSTSTA protofibril appears tightly restrained within the crystal lattice as evidenced by mean B-factors of 0.92 Å^2^. The modelled water molecule also appears well ordered, particularly in the structure of L-GSTSTA determined by MicroED where it has a B-factor 3.28 Å^2^. The single coordinated water is hydrogen bonded to serine 708, the C-terminus of a symmetry related strand, and the backbone of threonine 709 in the mating sheet (Fig. 4, S12, S13, and Table S4).

**Figure 4.**
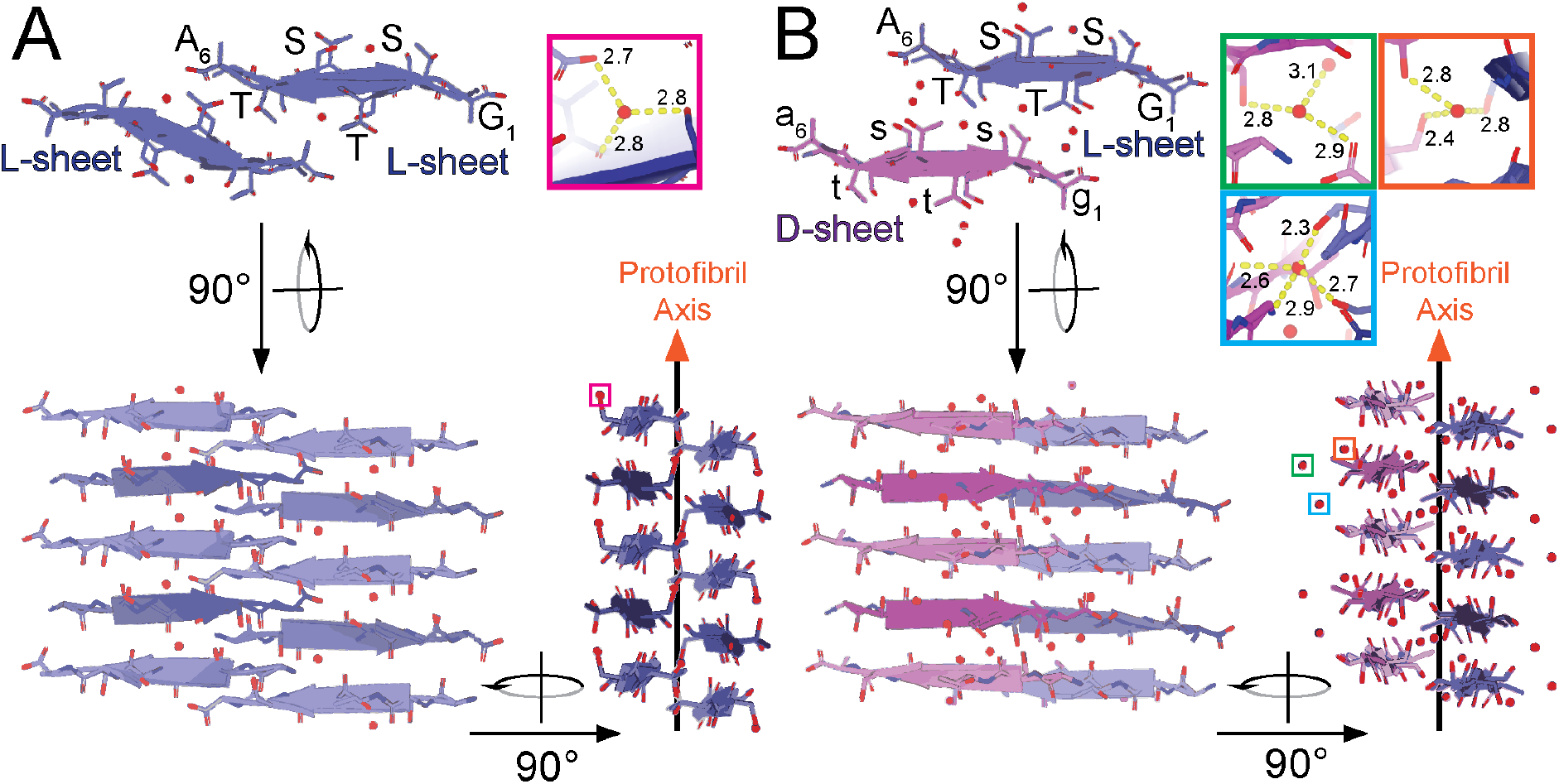
Views of protofibrils of L-GSTSTA (A) and racemic GSTSTA (B) represented by pair of sheets with a view down the protofibril axis; both structures derived by MicroED. A 90° rotation shows a side view of the protofibril with strands stacked along each sheet in an antiparallel fashion. Another 90° rotation shows a side view of the protofibril along the strand axis, showing a buckling of each sheet due to the tilting of strands away from or toward the protofibril axis. Chains colored such that blue represent L-peptides while magenta represents D-peptides. Lighter and darker shades of each color differentiate the orientation of strands within a sheet. Ordered waters found in each asymmetric unit are indicated by colored squares that correspond to insets of matching colors. Insets show magnified views of each water molecule with hydrogen bonds represented by the yellow dashed lines, labelled with their corresponding distances in Å.

### 2.7. Structure of racemic GSTSTA

In crystals of racemic GSTSTA, homochiral strands stack to form single enantiomer antiparallel beta sheets (Fig. 4B and Fig. S13). Like the homochiral L-GSTSTA sheets, racemic GSTSTA sheets are buckled with adjacent strands spaced 4.6 Å apart along each sheet (Fig. 4B). In the structure of racemic GSTSTA these sheets pack in alternating chirality, whereby each racemic GSTSTA protofibril is composed of one L-GSTSTA and one D-GSTSTA sheet (Fig. 4B). The packing of D-GSTSTA sheets against their L-GSTSTA mates in the racemic fibril differs from that seen in homochiral fibrils of L-GSTSTA. An alignment of the two protofibrils shows D-GSTSTA sheets displaced by approximately 5.3Å compared to their corresponding L counterparts in the homochiral fibril (Fig. 4 and Fig. S15). As a result of this displacement sheets are spaced farther apart (7-8Å) in the racemic GSTSTA protofibril (Fig. 4 and S12).

The longer spacing between sheets in the racemic GSTSTA protofibril is associated with bridging waters at its core (Fig. S12). These waters have extensive contacts along the protofibril, each hydrogen bonding with at least one threonine residue (Fig. 4, Fig. S13 and Table S4). Notably, the racemic GSTSTA structure shows a distinct rotamer for serine 710, which appears bound to an ordered water unlike its equivalent residue in the homochiral structure (Fig. 4, S15, Table S4). One water (water 1, Table S4) links serine 708 and threonine 711 on the same D sheet while also coordinating serine 708 of the adjacent L sheet. This water is isolated from the other waters found within the structure. A small network of waters near the protofibril core links the carboxylate of one strand to threonine 711 of a symmetry related strand (Fig. 4, Table S4). As with the structure of homochiral L-GSTSTA, peptide atoms and bound waters in racemic GSTSTA show low B-factors.

## 3. Discussion

Ice nucleation by *P. syringae* is linked to its expression of surface proteins including InaZ (Wolber *et al*., 1986). While full length InaZ and InaZ fragments help nucleate ice (Green & Warren, 1985; Kobashigawa Yoshihiro *et al*., 2005), individual InaZ repeats do not (Han *et al*., 2017). However, at the high concentrations required for crystallization, GSTSTA repeats self-assemble into a protofibrillar structure of corrugated beta sheets (Fig. S14). Similar structures are formed by both racemic GSTSTA and L-GSTSTA, and both contain ordered waters bridging tightly packed antiparallel beta sheets (Fig. 4, Fig. S12 and Fig. S13). These waters may play role in helping stabilize the GSTSTA protofibril or could act as bridges or templates for solvent ordering at low temperatures. While we have no evidence to suggest that GSTXT[A/S] repeats facilitate the formation of amyloid-like InaZ protofibrils, our structures of GSTSTA present an opportunity to analyze the interactions between polar residues in InaZ repeats and ordered solvent molecules at atomic resolution.

The structures of entantiomerically pure and racemic GSTSTA present a platform for comparison of homochiral and racemic amyloid protofibrils (Fig. S16). To evaluate the packing of each GSTSTA protofibril, we look at the categorization of strand packing in amyloid fibrils through homosteric zipper classes, first proposed by Sawaya and colleagues (Sawaya *et al*., 2007) and later by Stroud (Stroud, 2013). Many of these classes have been experimentally observed in amyloid crystals (Nelson *et al*., 2005; Sawaya *et al*., 2007). Homochiral GSTSTA forms a class-8 zipper in which two in register, antiparallel beta sheets meet, related by a 180-degree rotation normal to the protofibril growth axis (Sawaya *et al*., 2007; Stroud, 2013). The racemic GSTSTA structure resembles a class-8 zipper but is distinct in that two sheets of opposite handedness come together to form the protofibril (Fig. S16). Because of this similarity to a class-8 zipper, we label this arrangement class-8bar (Fig. S16).

The increased propensity for crystallization by racemic mixtures could be exploited to facilitate growth of amyloid crystals. The symmetry present in racemic amyloid crystals would have to accommodate the packing of homochiral protofibrils into the racemic lattice or allow the formation of racemic protofibrils (Yeates & Kent, 2012), as is the case with GSTSTA. Our experiments in high-throughput crystallographic trials of GSTSTA confirm the expected higher propensity for crystallization by racemic mixtures (Yeates & Kent, 2012) (Fig. S3) yielding a high number of conditions that contain sub-micron sized crystals suitable for MicroED. The facile determination of *ab initio* structures from these crystals demonstrates how MicroED combined with solid phase polypeptide synthesis (Dawson *et al*., 1994; Merrifield, 1986) can expand the reach of racemic crystallography to sub-micron sized crystals.

## 4. Methods

### 4.1. Sequence analysis of ice nucleation proteins

The sequence of the ice nucleating protein InaZ from *Pseudomonas syringae* was screened for the existence of hexameric degenerate repeat motifs that contained one or more threonine residues (Fig. S1). Repeats were then evaluated by their propensity for amyloid fibril formation by ZipperDB (Fig. S1). For each, a Rosetta energy score was calculated. A single repeat, GSTSTA, was chosen from this list of hexameric sequences. This segment appears five times identically across the sequence of InaZ, first at residue 707, and is part of a group with the consensus motif GSTXT[A/S] that appears 59 times across the InaZ sequence.

### 4.2. Synthesis, purification, and characterization and crystallization of L- and D-enantiomers of the InaZ derived peptide GSTSTA

The L-enantiomer of GSTSTA was purchased from Genscript with 98% purity. The D-enantiomer of GSTSTA was synthesized by solid phase peptide synthesis and purified using a Waters Breeze 2 HPLC System in reversed phase, buffered with trifluoroacetic acid (Fig. S2). The two enantiomers were qualified by ESI-MS on a Waters LCT Premier. The L-enantiomer spectrum showed a [M+H]^+^ peak of 523.30 g/mol (expected 523.22) and a dimer [M+M+H]^+^ peak of 1045.6 g/mol (expected 1045.44). The D-enantiomer spectrum showed a [M+H]^+^ peak of 523.24 g/mol (expected 523.22) and a dimer [M+M+H]^+^ peak of 1045.49 g/mol (expected 1045.44) (Fig. S2).

Crystals of L-GSTSTA were grown as follows: lyophilized peptide was weighed and dissolved in ultrapure water at concentrations between 82 and 287 mM, assuming a 1:1 ratio of peptide to trifluoroacetic acid (TFA) in the lyophilized powder. Crystals were grown at room temperature by the hanging-drop method in a high-content 96-well Wizard screen. Out of the many crystallization trials that yielded crystals, those in the condition containing, 0.1M CHES buffer, pH9.5, and 10% (w/v) PEG 3000, were used for microfocus x-ray data collection. Another promising condition was optimized by the hanging-drop method in 24-well plates. That condition contained 0.1M McIlvaine (citrate-phosphate) buffer, pH 4.2, and 12.5% (w/v) of PEG 8000 and 0.1M sodium chloride and was used to grow crystals of L-GSTSTA in batch.

Crystals of racemic GSTSTA were grown as follows: lyophilized powders of L-GSTSTA and D-GSTSTA were separately weighed and dissolved in ultrapure water so that the concentration of the two enantiomers matched. Crystal formation was screened at concentrations ranging from 82 to 123 mM after accounting for TFA. Control trays containing only L- or D-GSTSTA were prepared simultaneously alongside racemic screens. All three trays were stored and monitored at room temperature with crystal formation observed in various conditions. Images of every well were collected after 3 hours, 1 day, 3 days, 5 days, and 7 days and crystal formation monitored over time. A condition containing 0.1M imidazole, pH 8.0, and 10% (w/v) PEG 8000 produced the best crystals.

Crystals were grown in batch for data collection by MicroED. Lyophilized L-GSTSTA peptide was weighed and dissolved in 0.1M McIlvaine (citrate-phosphate) buffer, pH 4.2, and 12.5% (w/v) of PEG 8000 and 0.1M sodium chloride to an effective final concentration of 123 mM, mimicking the final concentration of a hanging drop in the 24-well optimization. Lastly, the solution was seeded with crystal needles extracted from crystals grown in the 24-well optimization described above. Batch crystals of racemic GSTSTA were grown from lyophilized L-GSTSTA and D-GSTSTA that were separately weighed and dissolved in 0.1M imidazole buffer, pH 8.0, containing 10% (w/v) of PEG 8000 to a final concentration for each enantiomer of 50 mM after accounting for the mass contributed by TFA.

### 4.3. Microfocus x-ray data collection and structure determination

Crystals of L-GSTSTA were harvested from a 96-well hanging drop using MiTeGen loops and flash frozen in liquid nitrogen. No additional cryoprotectant was used other than the PEG 3000 already present in the mother liquor. 72 diffraction images were collected with an oscillation range of 5° from a single crystal; 40 of these were indexed and integrated. Crystals of racemic GSTSTA were harvested from a 96-well hanging drop using MiTeGen loops and flash frozen in liquid nitrogen. No additional cryoprotectant was used other than the PEG 8000 already present in the buffer. 144 diffraction images were collected with an oscillation range of 2.5° from a single crystal; 64 of these were indexed and integrated.

Both homochiral and racemic GSTSTA crystals were diffracted under cryogenic conditions (100K) at the Advanced Photon Source (APS) beamline 24-ID-E, equipped with an ADSC Q315 CCD detector, using 5-μm beam with a 0.979-Å wavelength. Signal was limited only by the edge of our detector at approximately 1.1Å; as such, perhaps higher resolution data could be achieved by modifying our experimental geometry and/or adjusting the energy of the x-ray beam in our experiment. Data from both homochiral and racemic crystals were reduced in XDS (Kabsch, 2010), yielded *ab initio* solutions by SHELXT and SHELXD (Sheldrick, 2008). The phases obtained from these coordinates produced maps of sufficient quality for subsequent model building in Coot (Emsley *et al*., 2010). The resulting models were refined against the measured data using Phenix(Adams *et al*., 2010).

### 4.4. Electron microscopy, MicroED data collection and structure determination

Crystals were prepared for MicroED data collection following a variation of procedures detailed in Rodriguez *et al*. (Rodriguez *et al*., 2015) as follows: Following a 1:2 dilution in ultrapure water, crystals were applied to glow discharged grids of type (Pelco easiglow) 300 mesh Cu 1/4. Excess liquid was blotted off onto filter paper wet with 4-μL of ultrapure water to avoid salt crystal formation. Grids were plunge frozen into liquid ethane using a vitrobot (FEI). Grids were then initially stored in liquid nitrogen before being transferred to a liquid-nitrogen-cooled Gatan 626 cryo-holder for insertion and manipulation within the electron microscope.

MicroED data were collected from three sub-micron-thick needle crystals of L-GSTSTA and two sub-micron-thick needle crystals of racemic GSTSTA. Briefly, frozen hydrated crystals of either L-GSTSTA and racemic GSTSTA were visually inspected in overfocused diffraction mode on a cryo-cooled FEI Tecnai F20 microscope operated at 200 kV (Janelia Research Campus). Diffraction patterns used for structure determination were collected on a TVIPS TemCam-F416 CMOS detector in rolling-shutter mode. For L-GSTSTA, diffraction patterns were collected during unidirectional rotation with two second exposures. For racemic GSTSTA, diffraction patterns were collected during unidirectional rotation with three second exposures. A rotation rate of 0.30° s^-1^ and rotation angles ranging from −63° to 72° were used for both. Beam intensity was held constant, with an average dose rate of 0.003-0.005 e^-^ Å^-1^ sec^-1^ or ~0.01 e^-^ Å^-2^ per image, corresponding to a total dose of ~1-3 e^-^ Å^-2^ per dataset. Data were recorded at an effective camera length of 730 mm, the equivalent of a sample to detector distance of 1156 mm in a corresponding lensless system. All diffraction was performed using a circular selected area aperture of ~1μm^2^ in projection.

Diffraction movies were converted to the SMV file format using TVIPS tools as previously described (Hattne *et al*., 2015). Indexing and integration were performed in XDS. Partial datasets from three L-GSTSTA crystals were sorted and merged in XSCALE. In total, intensities from 196 diffraction images were merged. An *ab initio* solution was achieved, using SHELXD (Sheldrick, 2008). To achieve a complete data set from racemic GSTSTA crystals, the integrated diffraction intensities from partial datasets of two different crystals were sorted and merged in XSCALE. In total, intensities from 145 diffraction images were merged. An *ab initio* solution was achieved, using SHELXD and SHELXT (Sheldrick, 2008). Although XDS accurately differentiated the Laue classification for the racemic GSTSTA data, SHELXT, which does not rely on user input for space group selection, ensured a correct solution for the racemic data. SHELXT selected P21/C as the racemic space group, a choice corroborated by the systematic absences present in the data. The phases obtained from L-GSTSTA and racemic GSTSTA coordinates produced by SHELX were used to generate maps of sufficient quality for subsequent model building in Coot (Emsley *et al*., 2010). The resulting models were refined with Phenix (Adams *et al*., 2010), using electron scattering form factors, against the measured data.

### 4.5. Analysis of homochiral and racemic GSTSTA structures

In the analysis of hydrogen bonding and assembly interactions of each L-GSTSTA structure, an assembly of 4 strands, composed of 2 pairs in mating sheets, was used to find all unique hydrogen bonds, while racemic GSTSTA required an assembly of 12 strands, composed of 3 strands from a pair of mating sheets and 6 more strands related by translation along the protofibril axis to achieve a unique set of hydrogen bonds. Hydrogen bonds were tabulated from this structure using the program Hbplus (McDonald & Thornton, 1994).

Distances between strands along a sheet were calculated as differences between alpha carbons of one strand with its neighbor along the same sheet. These distances were calculated for both GSTSTA and GNNQQNY using PDB ID 1YJP(Sawaya *et al*., 2016). The angle between a strand and its corresponding sheet was calculated against the plane formed by alpha carbons along that sheet.

### 4.6. Analysis of phases in structures determined by MicroED and x-ray crystallography

To analyze the distribution of phases associated with reflections measured from racemic crystals by both x-ray and electron diffraction, data reduction was performed in space group 1 (P1) and refined in Phenix against a model encompassing the entire unit cell of 4 strands. This model was obtained by applying all symmetry operations on the asymmetric unit of the P21/C structure. The refinement in P1 allowed symmetry to be broken, no longer restricting phases to 0 or 180 degrees as the phases changed in the case where coordinates deviated from their symmetry related positions. The resulting set of reflections and phases were analyzed in Matlab^®^. We plotted the observed and calculated magnitudes of each reflection against each other and the set fitted by linear regression. For each measured magnitude, the phases associated were plotted and showed a bimodal distribution. Histograms were drawn using these data to evaluate phase distributions; the standard deviation of these was computed by merging the distributions around 0 and 180 degrees using a modulo operation.

### 4.7. Analysis of paired reflections in MicroED and x-ray crystallographic data

Merged data sets collected by either MicroED or microfocal x-ray crystallography were paired for homochiral and racemic crystals of GSTSTA. MicroED data mtz files were scaled against their corresponding x-ray counterpart where corresponding reflections were paired and missing reflections ignored within a single .mtz file. This was achieved using custom scripts and the Rstats program which scaled and compared common reflections between corresponding datasets. The corresponding distributions of Fourier magnitudes were then analysed using Matlab^®^ in which a best fit line was determined for each of the paired datasets. Zones were visualized using the HKL View program in which either h, k, or l were selectively set to zero.

## Acknowledgements

This project was inspired by conversations on racemic crystallography with Todd Yeates (UCLA). We thank Dan Shi (HHMI, Janelia Research Campus), David Boyer and Daniel Anderson (UCLA) for their assistance in data collection, Johan Hattne (UCLA) for helpful discussions and Janak Dadhaniya for assistance with figures.

## Supporting information

**Table S1.**
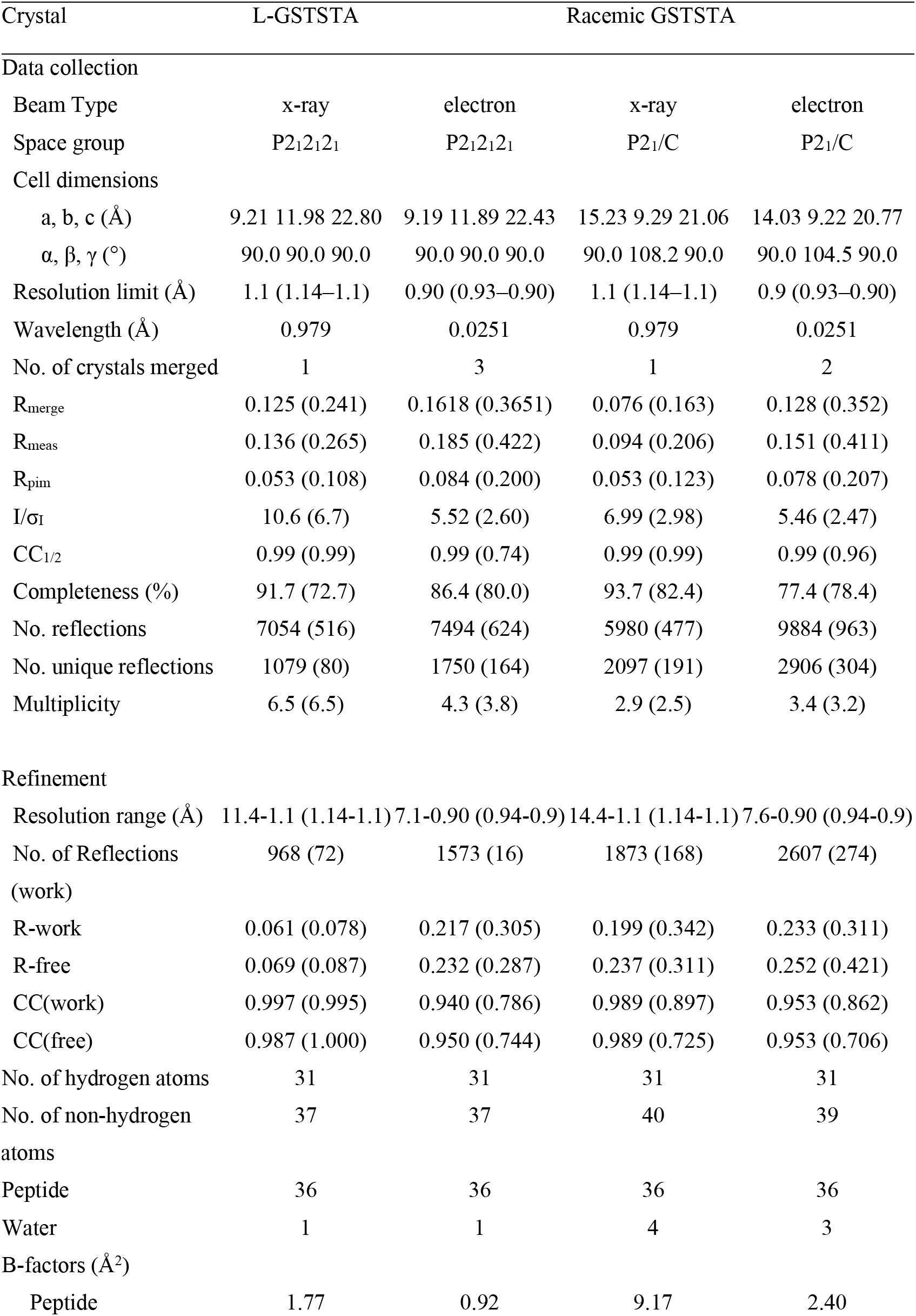
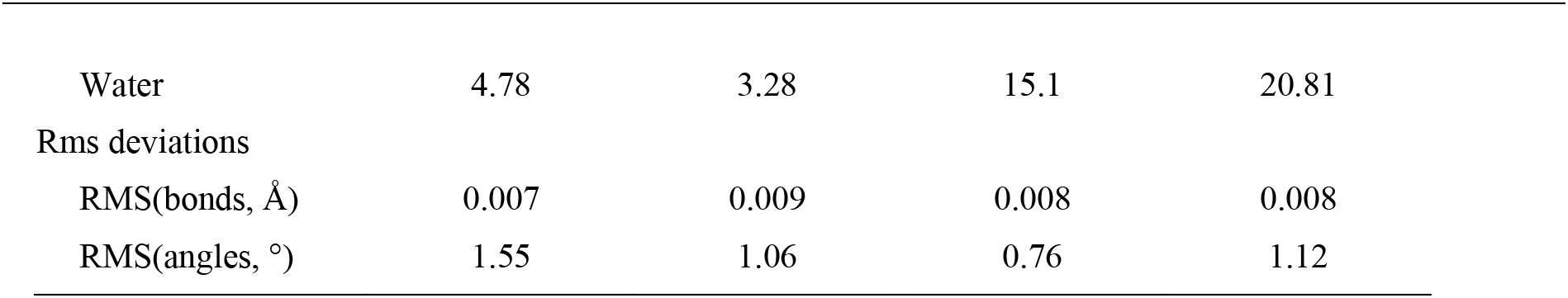
Crystallographic data collection and refinement statistics. Values in parentheses are for the highest-resolution shell. All modelled waters have an occupancy of 1 after refinement.

**Table S2.**
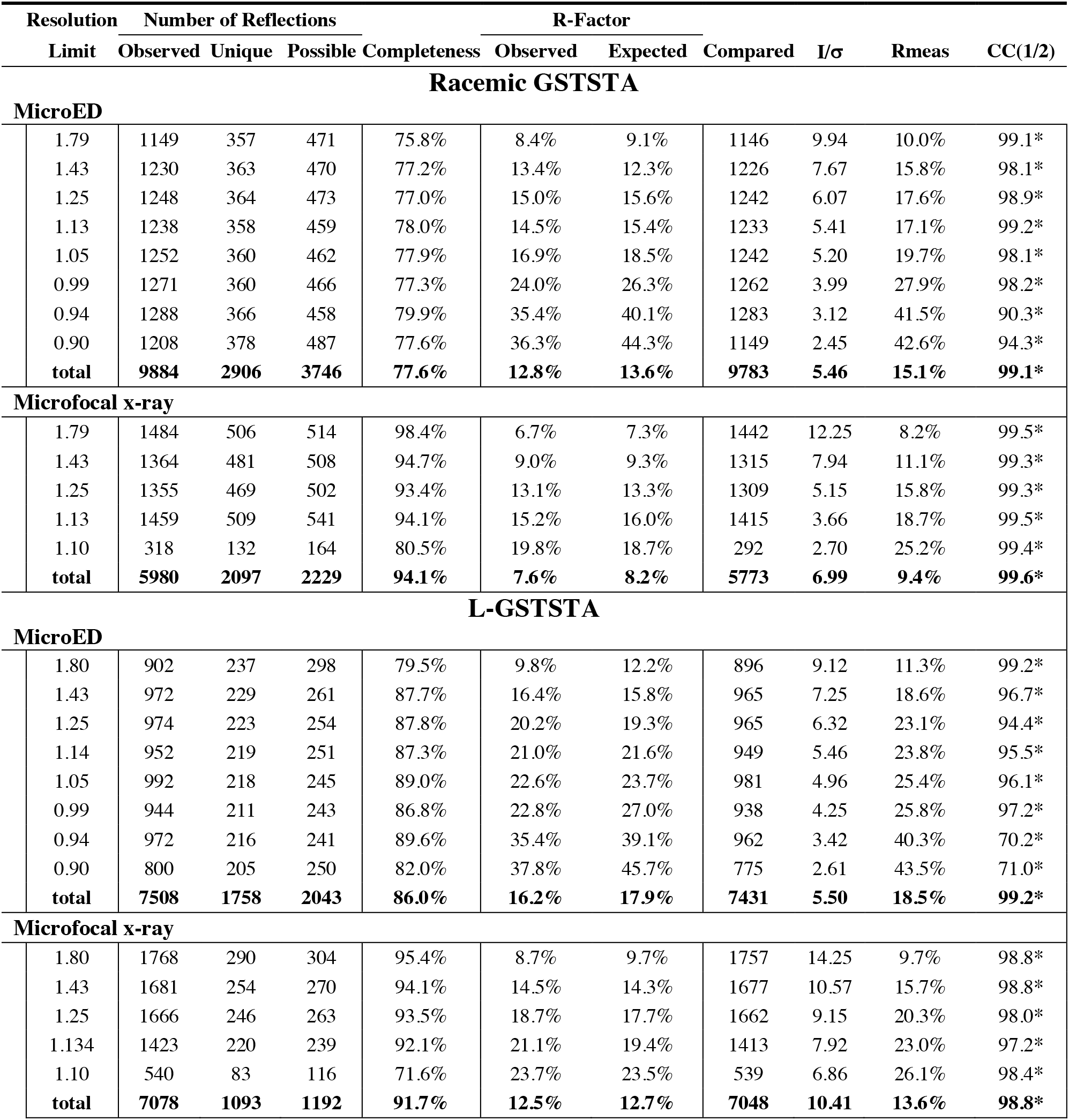
Data reduction statistics for homochiral and racemic GSTSTA crystals for microfocal x-ray diffraction and MicroED.

**Table S3.** Hydrogen bonding between adjacent strands in homochiral L-GSTSTA sheets. Peptide donor and acceptors are denoted as: chain name followed by a three-digit residue number and the associated three letter code. This table includes only unique hydrogen bonds between strands in a single beta sheet along the protofibril axis. All distances are measured in Ångströms, angles in degrees.

| **Donor** | **Atom** | **Acceptor** | **Atom** | **DA dist.** | **Type** | **CA to** | **DHA** | **H-A Dist.** | **H-A-AA** | **D-A-AA** |
| --- | --- | --- | --- | --- | --- | --- | --- | --- | --- | --- |
| B707-Gly | N | A712-Ala | O | 2.80 | MM | 5.1 | 165.5 | 1.80 | 139.8 | 144.7 |
| A712-Ala | N | B707-Gly | O | 2.95 | MM | 5.1 | 153.1 | 2.03 | 140.6 | 142.1 |
| B709-Thr | N | A710-Ser | O | 2.97 | MM | 5.2 | 154.6 | 2.04 | 139.6 | 146.2 |
| A710-Ser | N | B709-Thr | O | 2.82 | MM | 5.2 | 151.3 | 1.90 | 147.2 | 157.0 |
| A709-Thr | OG1 | B710-Ser | OG | 3.40 | SS | 4.1 | 153.7 | 2.48 | 94.0 | 99.2 |
| B711-Thr | N | A708-Ser | O | 2.89 | MM | 5.0 | 144.8 | 2.01 | 128.8 | 139.2 |
| A708-Ser | N | B711-Thr | O | 2.92 | MM | 5.0 | 154.7 | 1.98 | 137.6 | 138.0 |
| B711-Thr | OG1 | A708-Ser | OG | 2.68 | SS | 5.0 | 165.0 | 1.70 | 130.4 | 125.4 |

**Table S4.** Hydrogen bonds of modelled waters in homochiral and racemic GSTSTA structures. Peptide donor and acceptor names are abbreviated as the chain name followed by a three-digit residue number, a dash, and the associated three letter code. All hydrogen bonds in the table include a water as a donor or acceptor and are restricted to waters found in one asymmetric unit. All distances are measured in ångströms, angles in degrees. Only hydrogen bonds with distances below 3.2 Å are listed. The list includes potential hydrogen bonding partners, though not all might be satisfied at one time for a given atom.

| Donor Name | Atom Type | Acceptor Name | Atom Type | DA dist. | D-A-AA angle |
| --- | --- | --- | --- | --- | --- |
| **L- GSTSTA, X-Ray** |  |  |  |  |  |
| Water 1 | O | A709-Thr | O | 2.84 | 113.9 |
| Water 1 | O | C712-Ala | O | 2.78 | 109.1 |
| B708-Ser | OG1 | Water 1 | O | 2.72 |  |
| **Racemic GSTSTA, X-Ray** |  |  |  |  |  |
| Water 1 | O | A709-Thr | O | 2.93 | 121.1 |
| B708-Ser | OG1 | Water 1 | O | 2.82 |  |
| B710-Ser | OG1 | Water 1 | O | 2.85 |  |
| Water 2 | O | G712-D-Ala | O | 2.89 | 107.2 |
| Water 2 | O | Water 3 | O | 2.71 |  |
| E711-D-Thr | OG1 | Water 2 | O | 2.93 |  |
| Water 3 | O | Water 4 | O | 2.94 |  |
| Water 3 | O | E712-D-Ala | O | 2.86 | 102.6 |
| H707-Gly | N | Water 3 | O | 2.80 |  |
| C709-Thr | OG1 | Water 3 | O | 2.67 |  |
| Water 4 | O | C708-Ser | O | 2.78 | 122.3 |
| Water 4 | O | B711-Thr | O | 2.89 | 127.8 |
| H707-Gly | N | Water 4 | O | 2.90 |  |
| C709-Thr | OG1 | Water 4 | O | 3.02 |  |
| **L-GSTSTA, MicroED** |  |  |  |  |  |
| Water 1 | O | D712-Ala | O | 2.66 | 112.7 |
| Water 1 | O | B709-Thr | O | 2.76 | 113.3 |
| A708-Ser | OG1 | Water 1 | O | 2.76 |  |
| **Racemic GSTSTA, MicroED** |  |  |  |  |  |
| Water 1 | O | E711-D-Thr | OG1 | 2.81 | 126.9 |
| D708-D-Ser | OG1 | Water 1 | O | 2.42 |  |
| B708-Ser | OG1 | Water 1 | O | 2.76 |  |
| Water 2 | O | Water 3 | O | 3.05 |  |
| Water 2 | O | G712-D-Ala | O | 2.87 | 107.4 |
| E711-D-Thr | OG1 | Water 2 | O | 2.76 |  |
| Water 3 | O | H712-D-Ala | O | 2.62 | 108.1 |
| H707-Gly | N | Water 3 | O | 2.93 |  |
| C709-Thr | OG1 | Water 3 | O | 2.34 |  |
| B710-Ser | OG1 | Water 3 | O | 2.71 |  |

**Figure S1.**
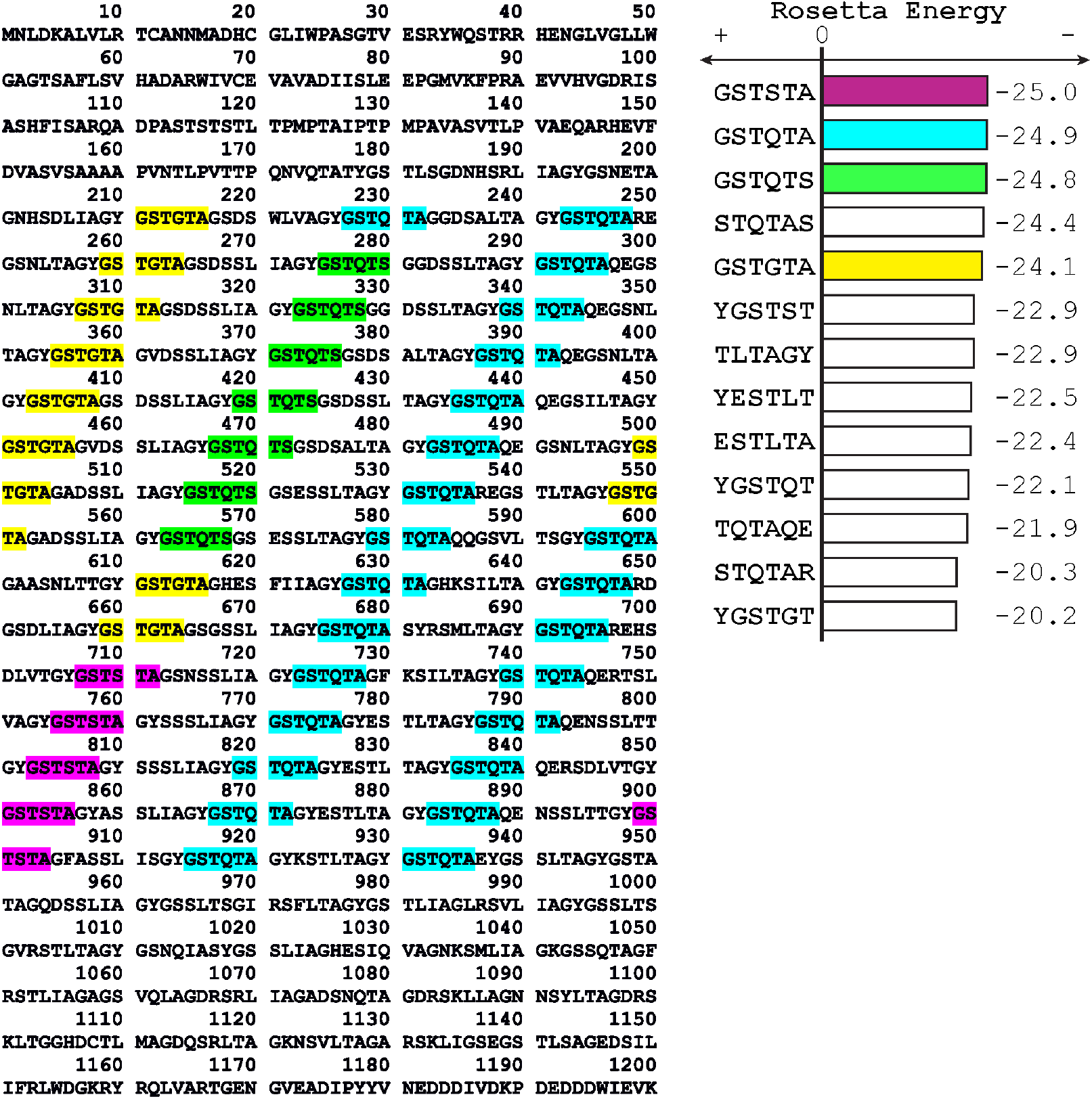
Sequence of the ice nucleation protein InaZ is shown with its degenerate hexameric repeats highlighted as follows: GSTGTA (yellow), GSTQTA (cyan), GSTSTA (magenta), and GSTQTS (green). The propensity for the hexamers to form steric zippers is shown on the left as Rosetta energy scores, determined by ZipperDB (Goldschmidt *et al*., 2010). This list of repeats is limited to those with Rosetta energy lower than −20 that containing at least two threonine residues and appear with frequency greater than or equal to five across the InaZ sequence.

**Figure S2.**
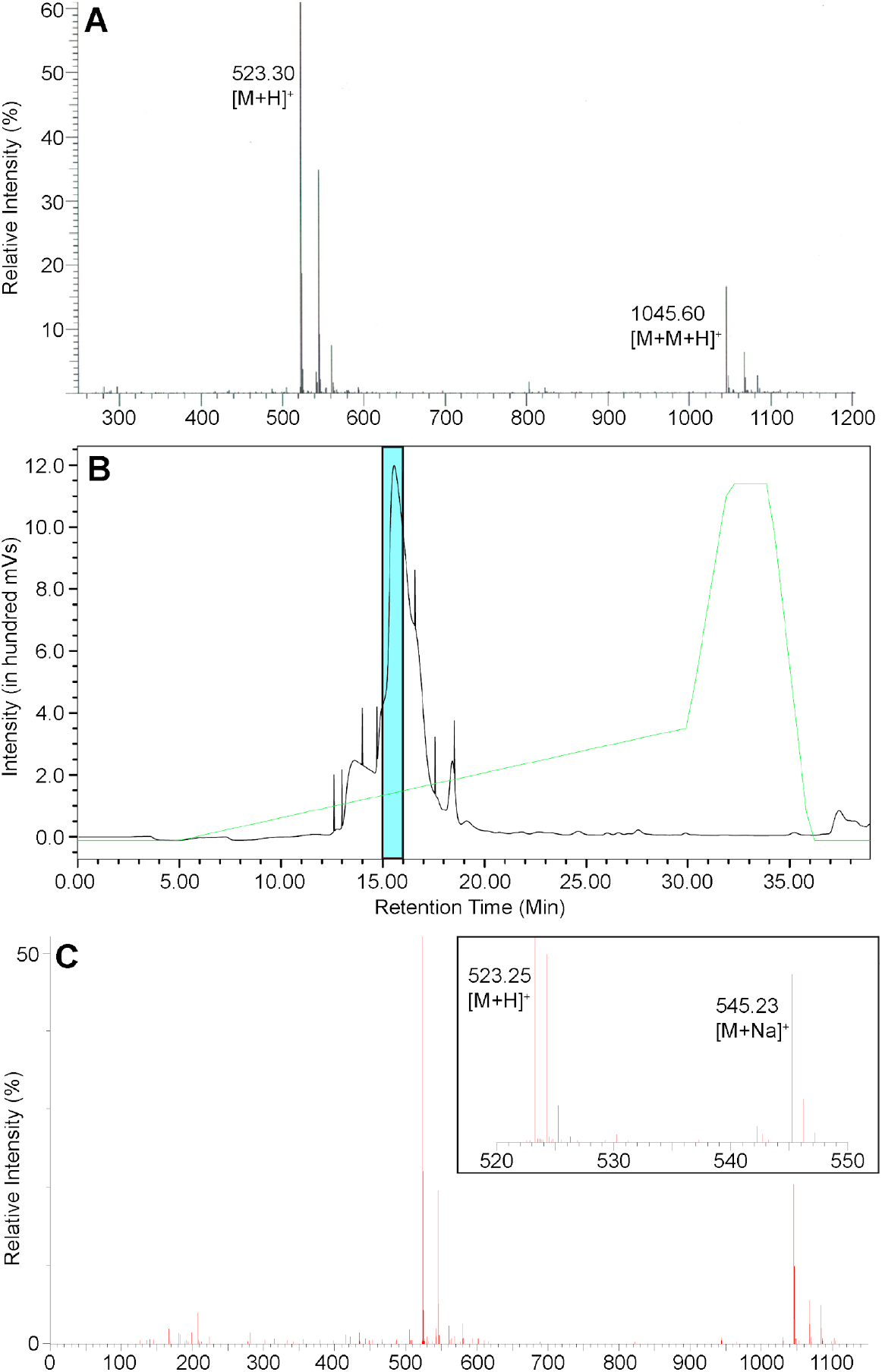
The mass trace of L-GSTSTA (A), purchased from Genscript, shows a [M+H]^+^ peak of 523.30 g/mol (expected 523.22) and a dimer [M+M+H]^+^ peak of 1045.6 g/mol (expected 1045.44). D-GSTSTA was synthesized and purified in-house by reverse-phase HPLC (B). The cyan shaded region highlights the HPLC fraction collected and lyophilized for crystallization experiments. The mass spectrum of D-GSTSTA (C) shows a [M+H]^+^ peak of 523.24 g/mol (expected 523.22) and a [M+Na]^+^ peak of 545.23 g/mol (expected 545.22).

**Figure S3.**
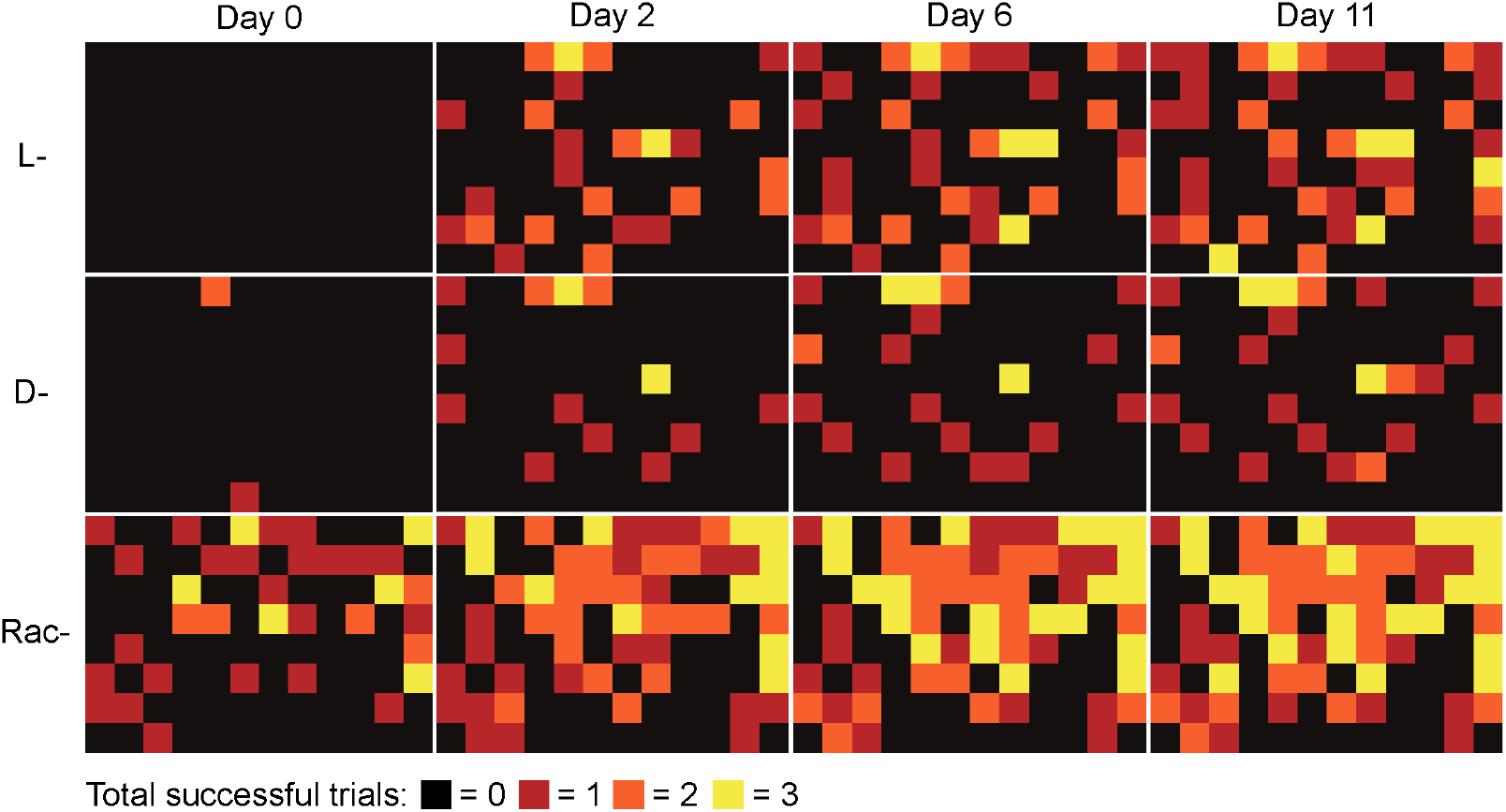
The number of successful crystallization trials in the 96-well crystal screens for L-GSTSTA, D-GSTSTA, and racemic GSTSTA were monitored and converted to heatmaps. Each well contained three hanging drops, each with different protein to buffer ratios. A count of crystals found in each of these three conditions was given a score from 0 to 3. No crystals in drops (black), one drop with crystals (red), two drops with crystals (orange), crystals in all three drops (yellow). The initial time point was three hours after setting up the screens (day 0), with data points collected up to 11 days post-setup.

**Figure S4.**
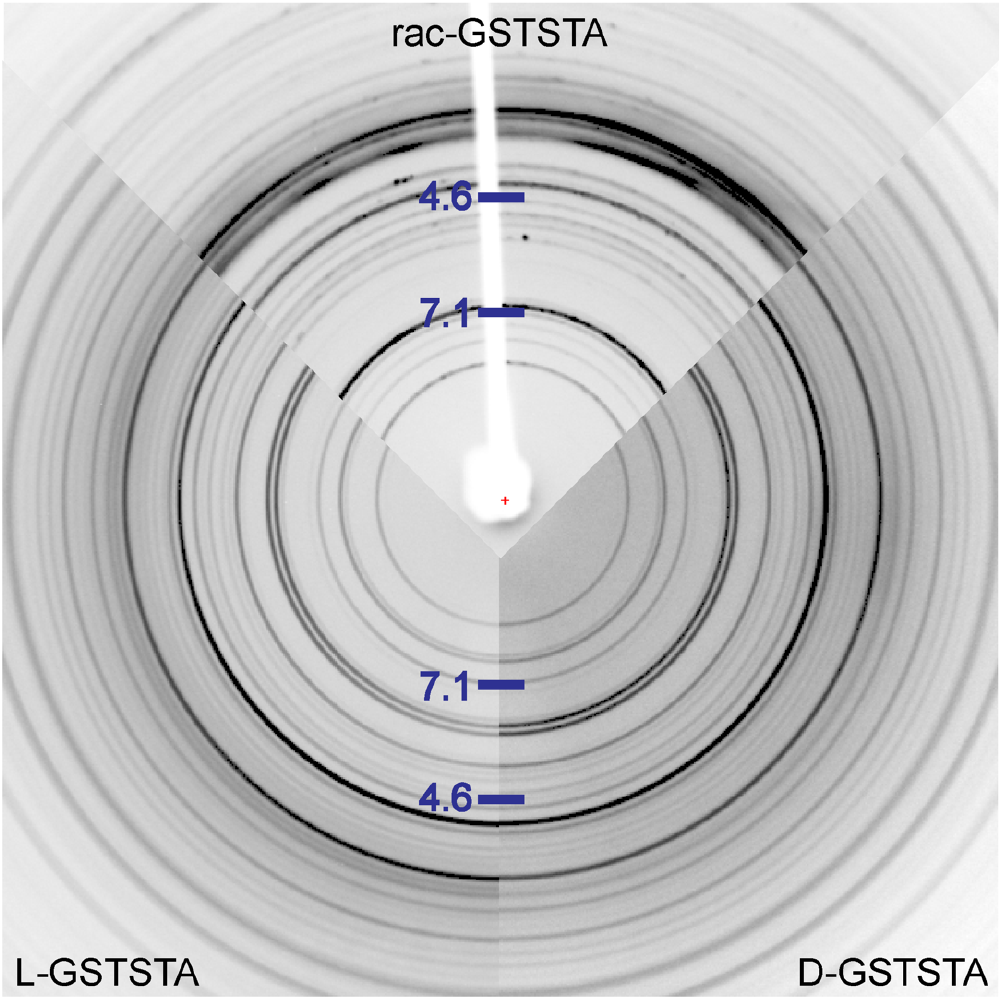
Comparison of powder diffraction patterns measured from L-GSTSTA (left), D-GSTSTA (right), and racemic GSTSTA (top) slurries show differences across all resolutions. Both patterns show faint rings at approximately 4.6Å, representing the approximate distance between strands along the fibril axis. The racemic pattern contains a prominent reflection at ~7.1Å, representative of overall sheet-to-sheet distances in its structure.

**Figure S5.**
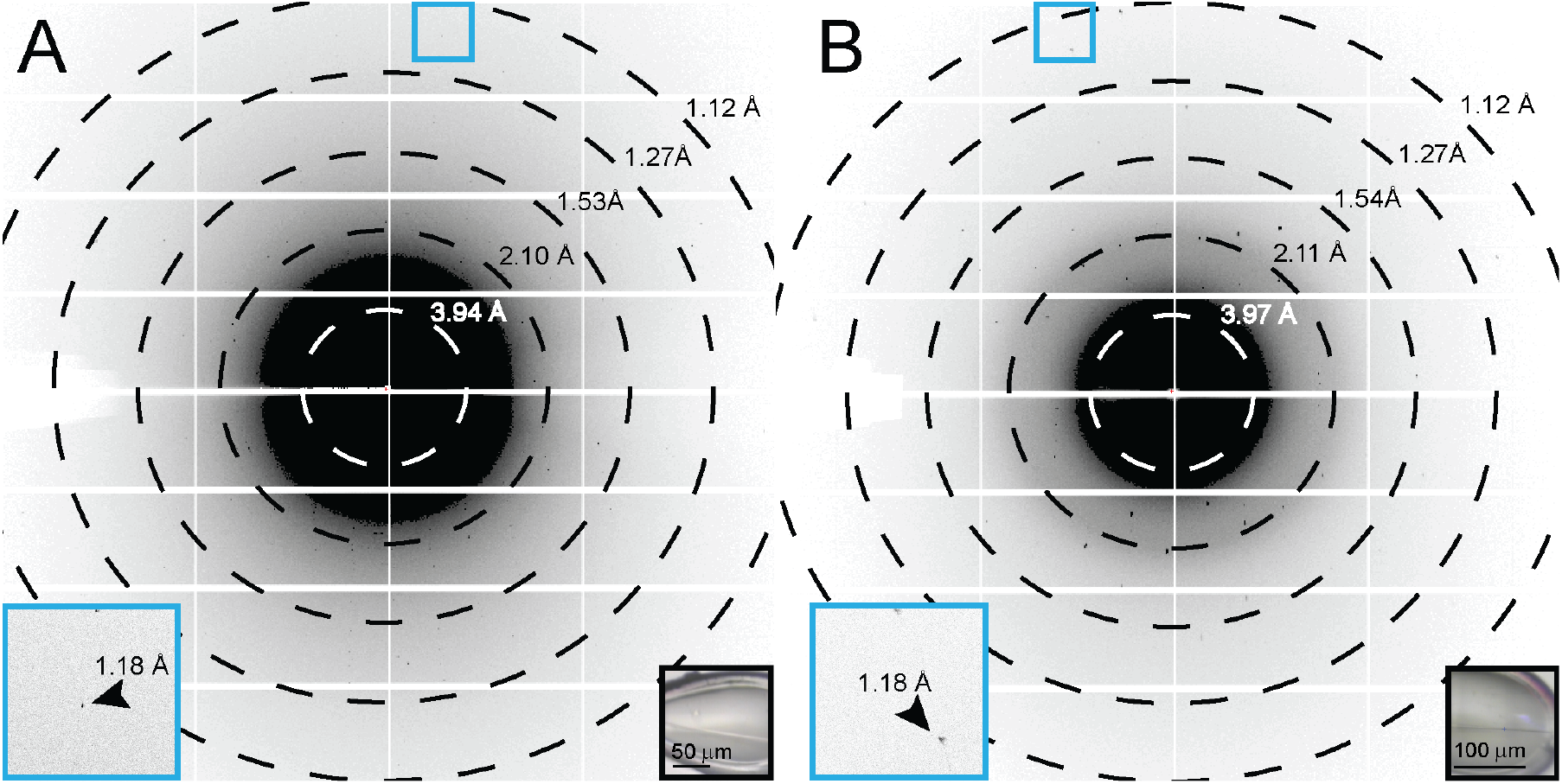
Single diffraction patterns of homochiral L-GSTSTA (A) or racemic GSTSTA (B) measured during continuous vector scanning, microfocus x-ray data collection. Each pattern corresponds to a 5° wedge (A) or 2.5° wedge (B) of reciprocal space. Black insets show in-line images of the crystals that were diffracted; blue squares correspond to magnified regions (blue insets) of the pattern that show diffraction near the detector edge at approximately 1.1Å resolution (black arrows). Resolution circles are indicated by rings; scale bars are 50 to 100 μm for L-GSTSTA and racemic GSTSTA, respectively.

**Figure S6.**
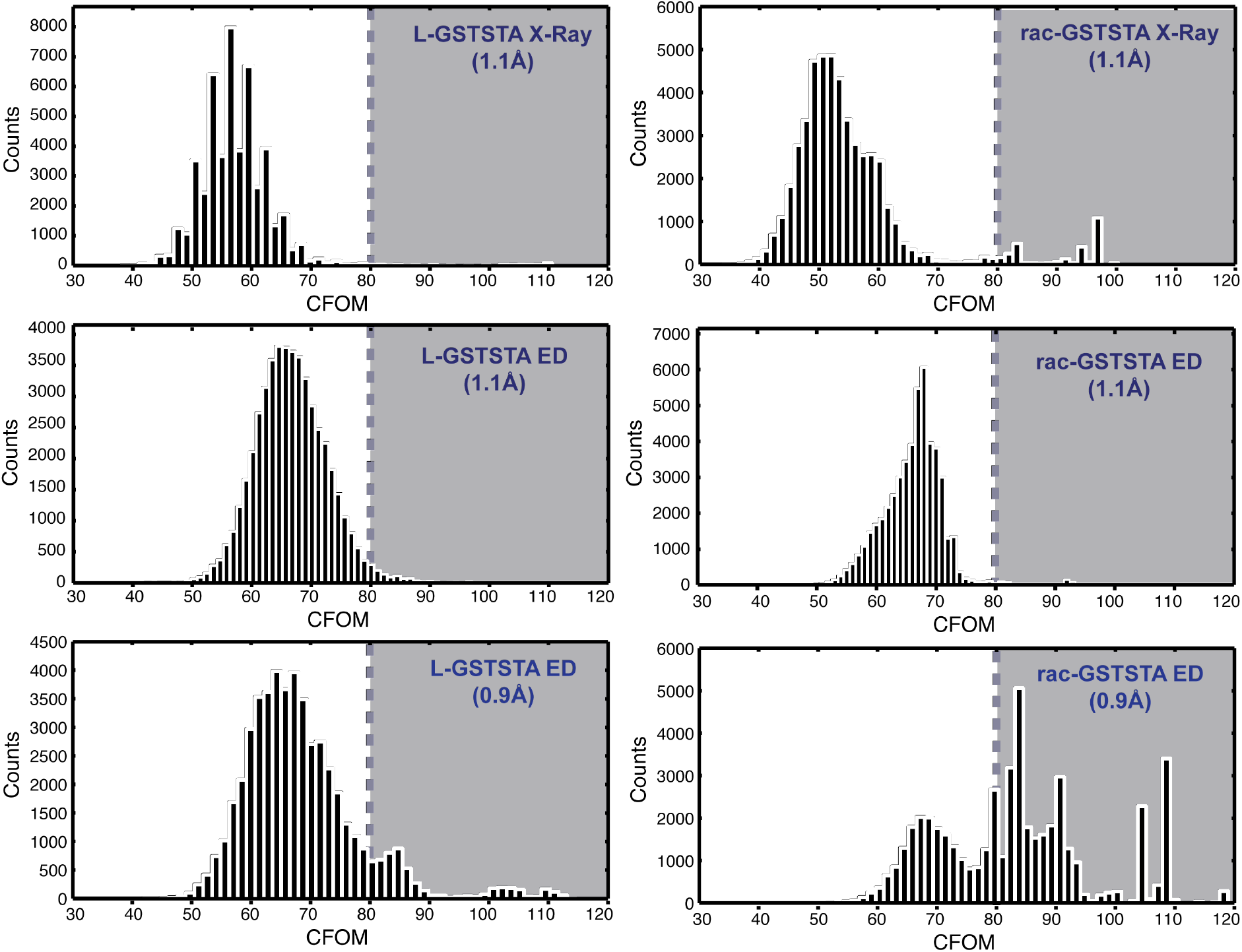
Histograms of the combined figure of merit (CFOM) scores for 50,000 trials by SHELXD indicate the approximate frequency of correct solutions. The shaded region in each plot, where CFOM scores are greater than 80, represents an area in which solutions have a high probability of being correct. Two sets of plots were generated from MicroED data: results of attempts using a truncated dataset that matches the resolution of the x-ray data (1.1Å) are shown in middle panels, while bottom panels show results of attempts using the measured resolution (0.9Å) for that data.

**Figure S7.**
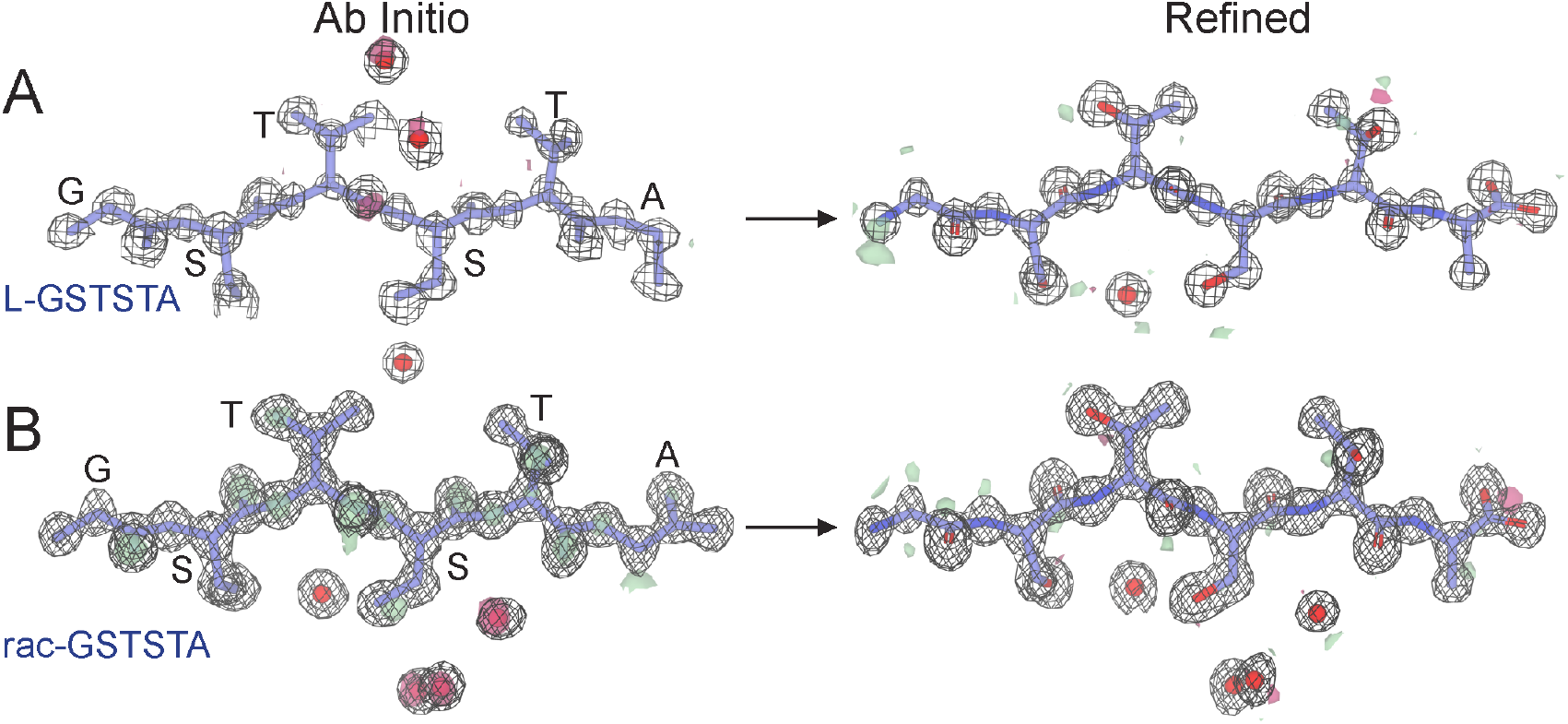
*Ab Initio* structures and electron density maps of L-GSTSTA (A) or racemic GSTSTA (B). Each map is in A is overlaid onto the initial atomic coordinates calculated by SHELXD from x-ray diffraction data. Each map in B is overlaid onto its corresponding refined model. The 2F_o_-F_c_ map represented by the black mesh is contoured at 1.2 σ. Green and red surfaces represent the F_o_-F_c_ maps contoured at 3.0 and −3.0 σ. Modelled waters are present as red spheres.

**Figure S8.**
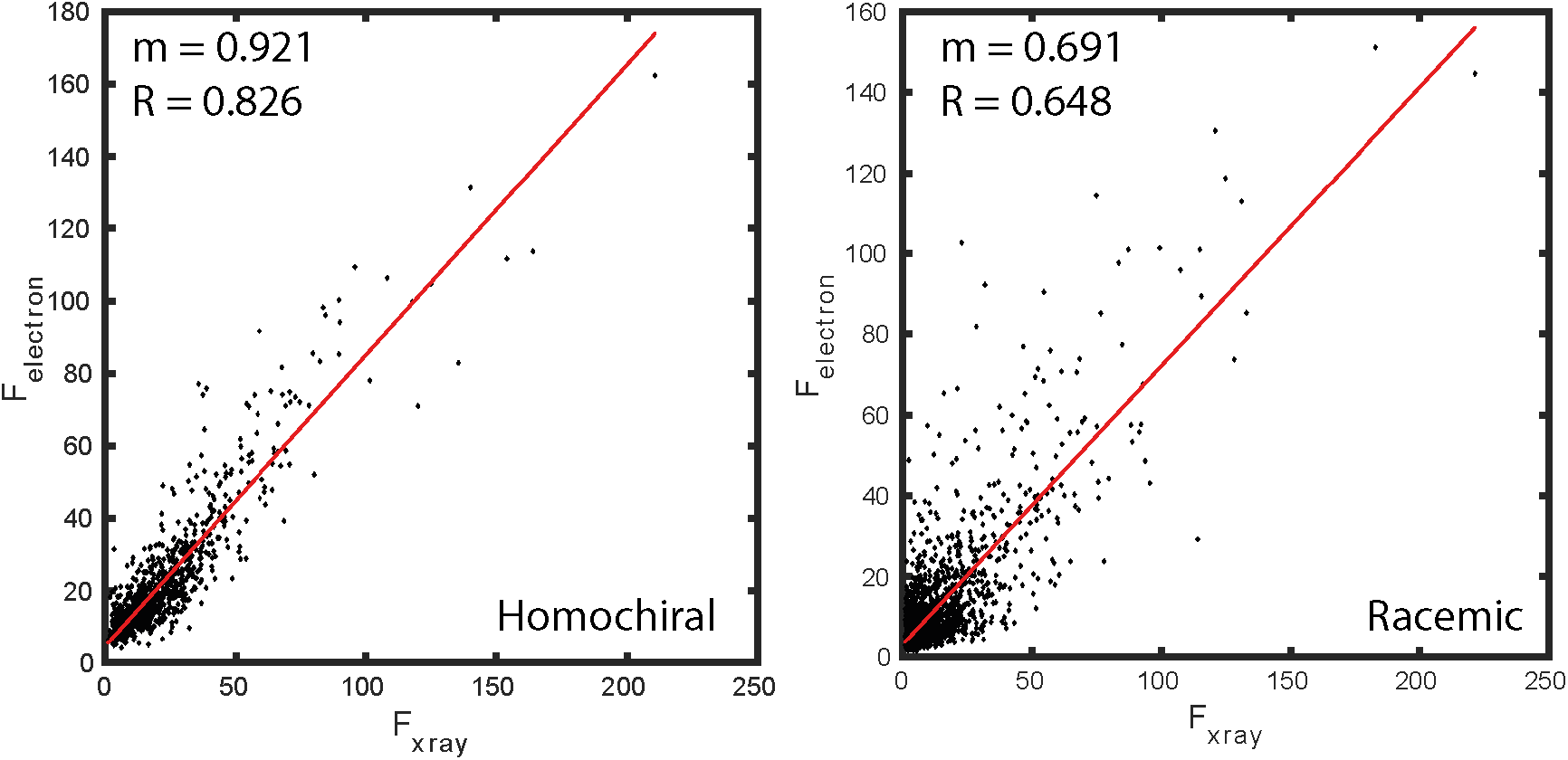
A plot of magnitudes (F) compared between reflections in datasets collected from homochiral crystals (left) or racemic crystals (right) by either electron or x-ray scattering shows a distribution that can be fit by linear regression, indicated by red lines with slope (m) and R-value.

**Figure S9.**
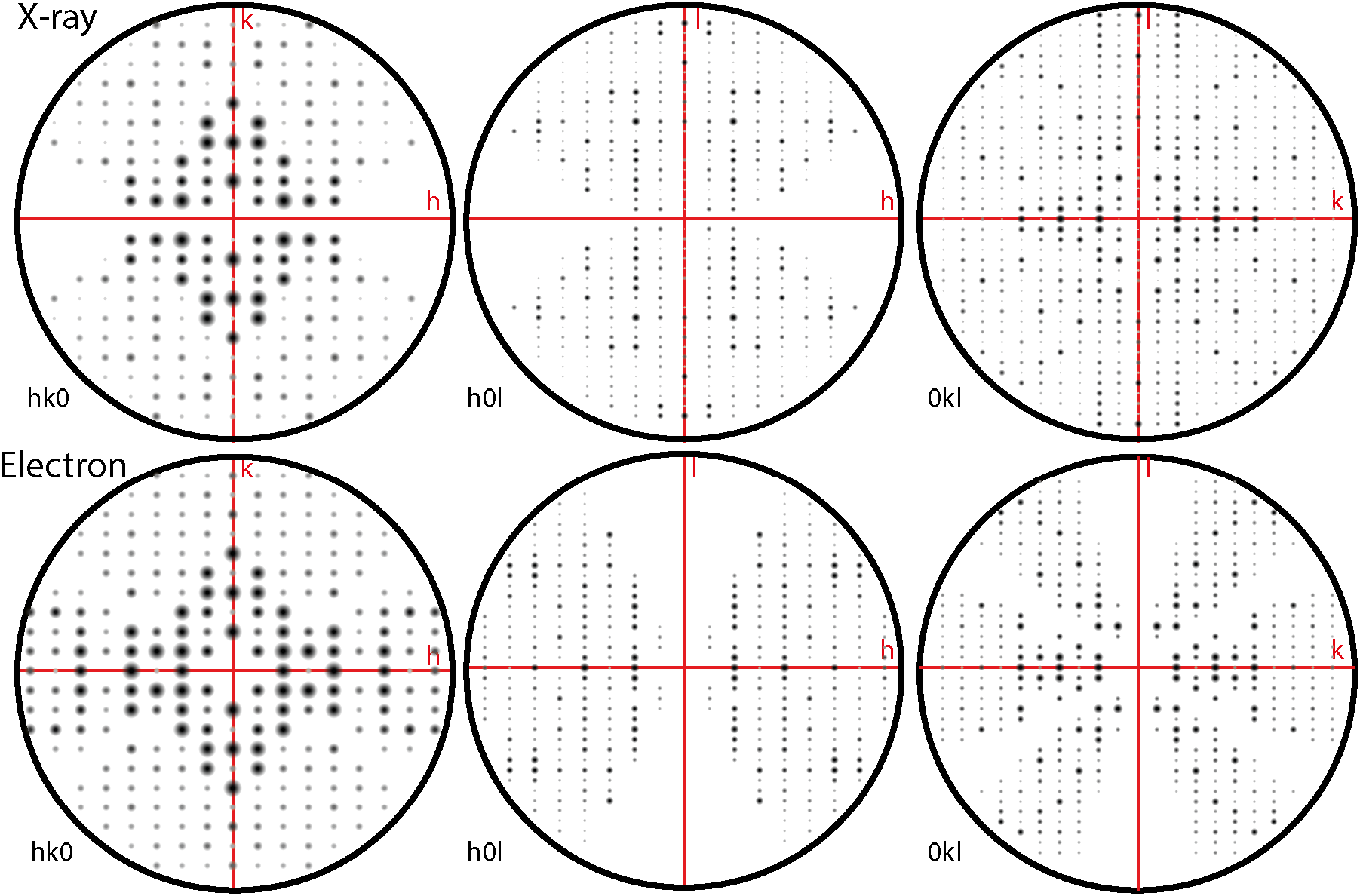
HKL Zone analysis for crystals of L-GSTSTA. Fourier magnitudes are shown as displayed by the HKL view software for reflections along principal zones of the reciprocal lattice. Zones where l=0 (left), k=0 (middle), h=0 (right) are shown for merged data collected by x-ray (top) and electron (bottom) diffraction. The black circle in each zone plot represents a resolution of 1.1Å; zone axes are labeled in red.

**Figure S10.**
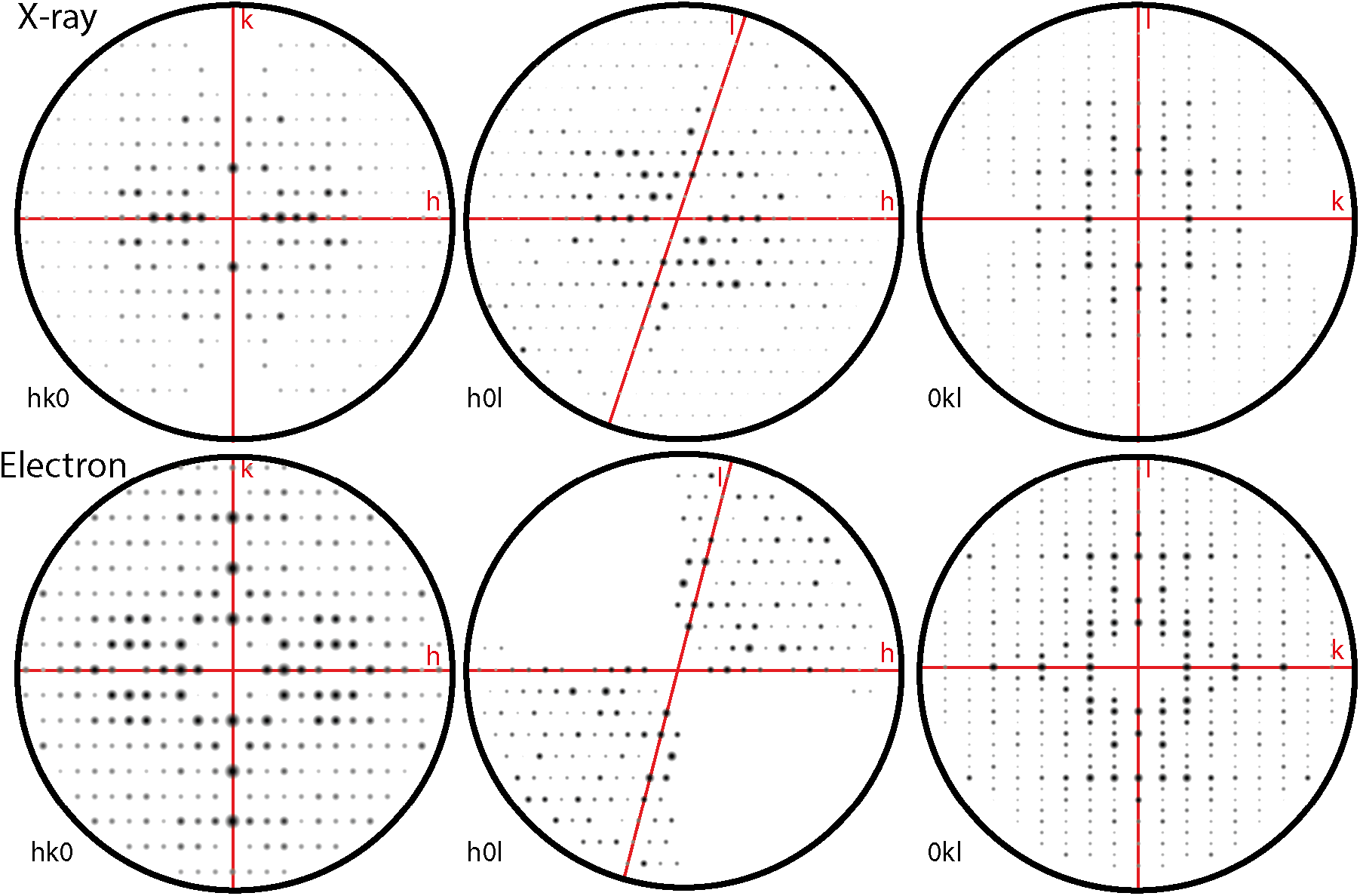
HKL Zone analysis for crystals of racemic GSTSTA. Fourier magnitudes are shown as displayed by the HKL view software for reflections along principal zones of the reciprocal lattice. Zones where l=0 (left), k=0 (middle), h=0 (right) are shown for merged data collected by x-ray (top) and electron (bottom) diffraction. The black circle in each zone plot represents a resolution of 1.1Å; zone axes are labeled in red.

**Figure S11.**
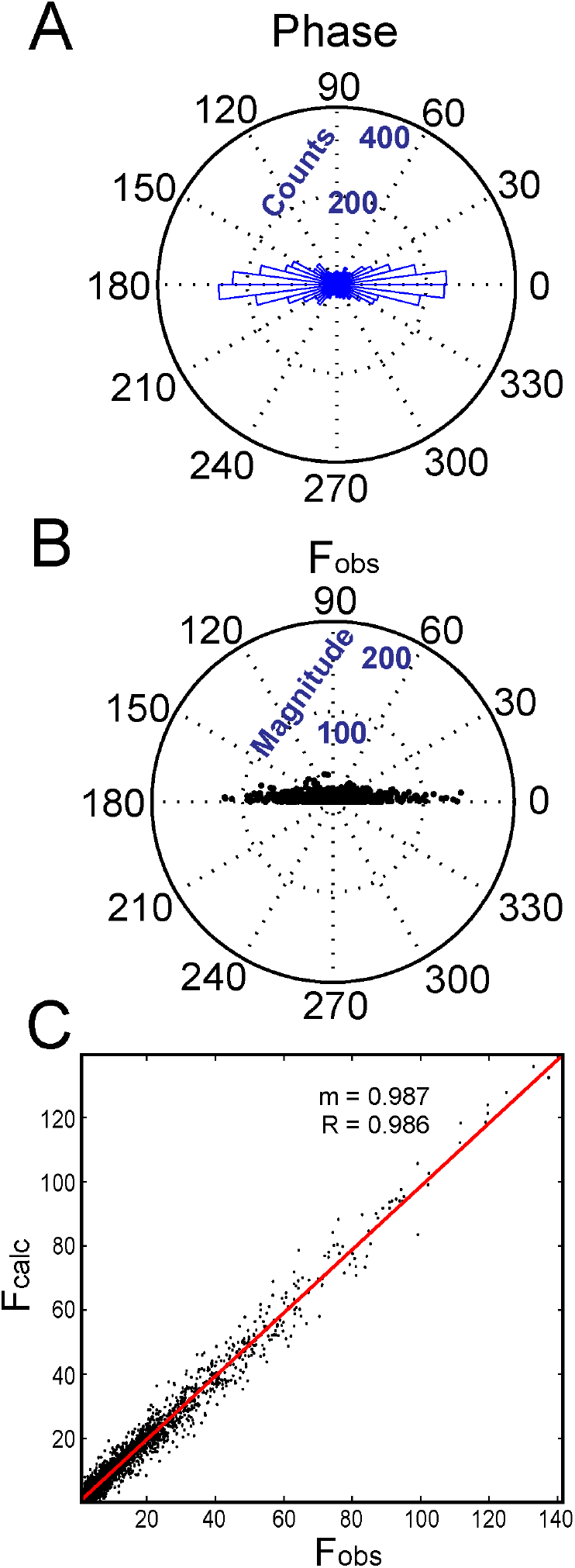
(A) The calculated phase associated with each reflection in the P1 refinement of racemic GSTSTA data obtained by x-ray diffraction was analyzed and plotted as a histogram along the unit circle. (B) The magnitude of each reflection is plotted as a function of the absolute value of its associated phase. (C) A plot of F_o_ vs. F_c_ values for each reflection in this data set shows a distribution that can be fit by linear regression, shown as a red line with slope m=0.987 and R-value 0.986.

**Figure S12.**
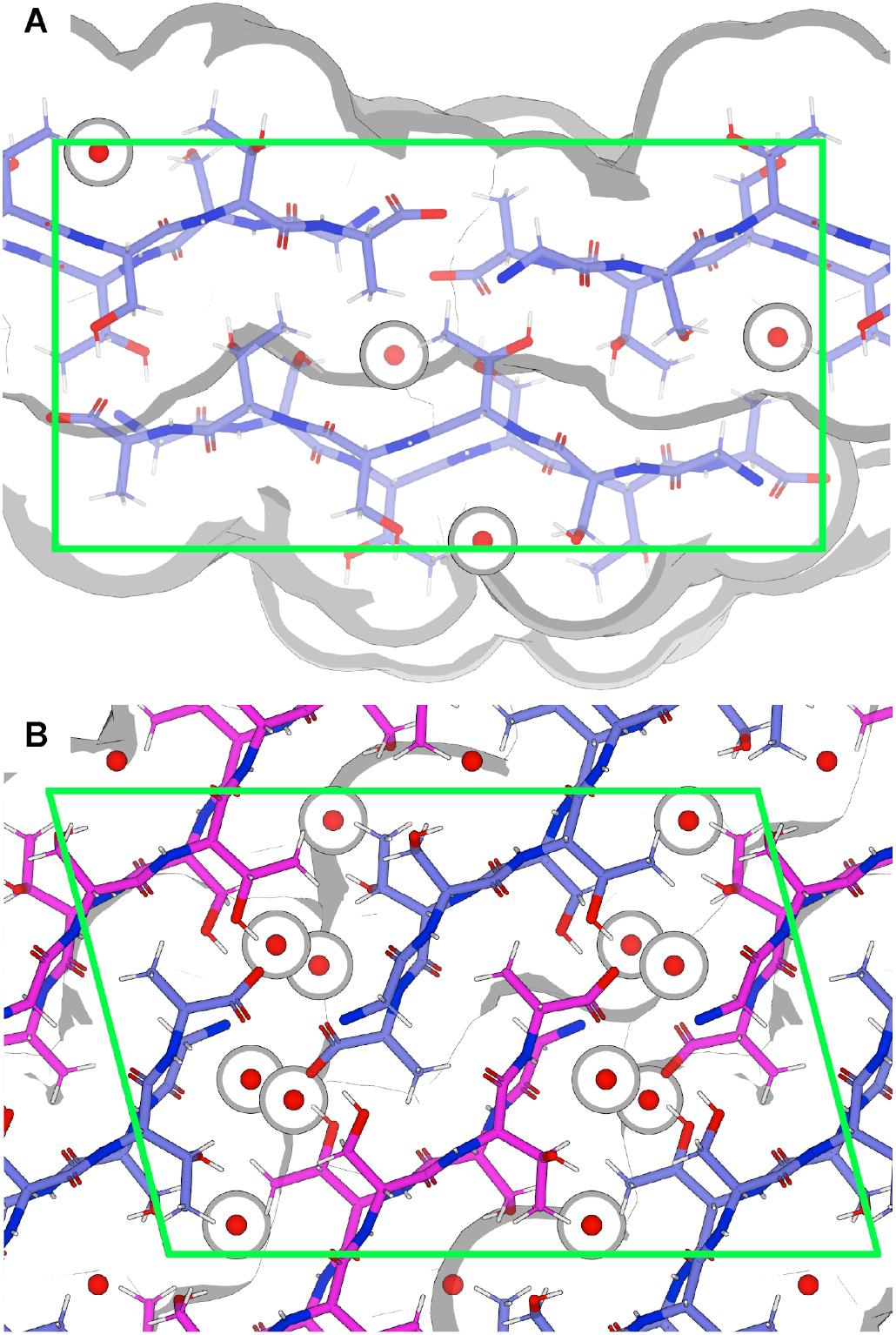
Views down the a-axis of L-GSTSTA (A) and racemic GSTSTA (B) structures are enclosed in green by their respective unit cells. L-GSTSTA strands are colored blue while D-GSTSTA strands are colored magenta. The volumes occupied by each structure are shown in white with edges defined by the shaded grey regions. Space-fill models represent the solvent accessible surface; ordered waters are represented by van der Waals radii of 1.4Å.

**Figure S13.**
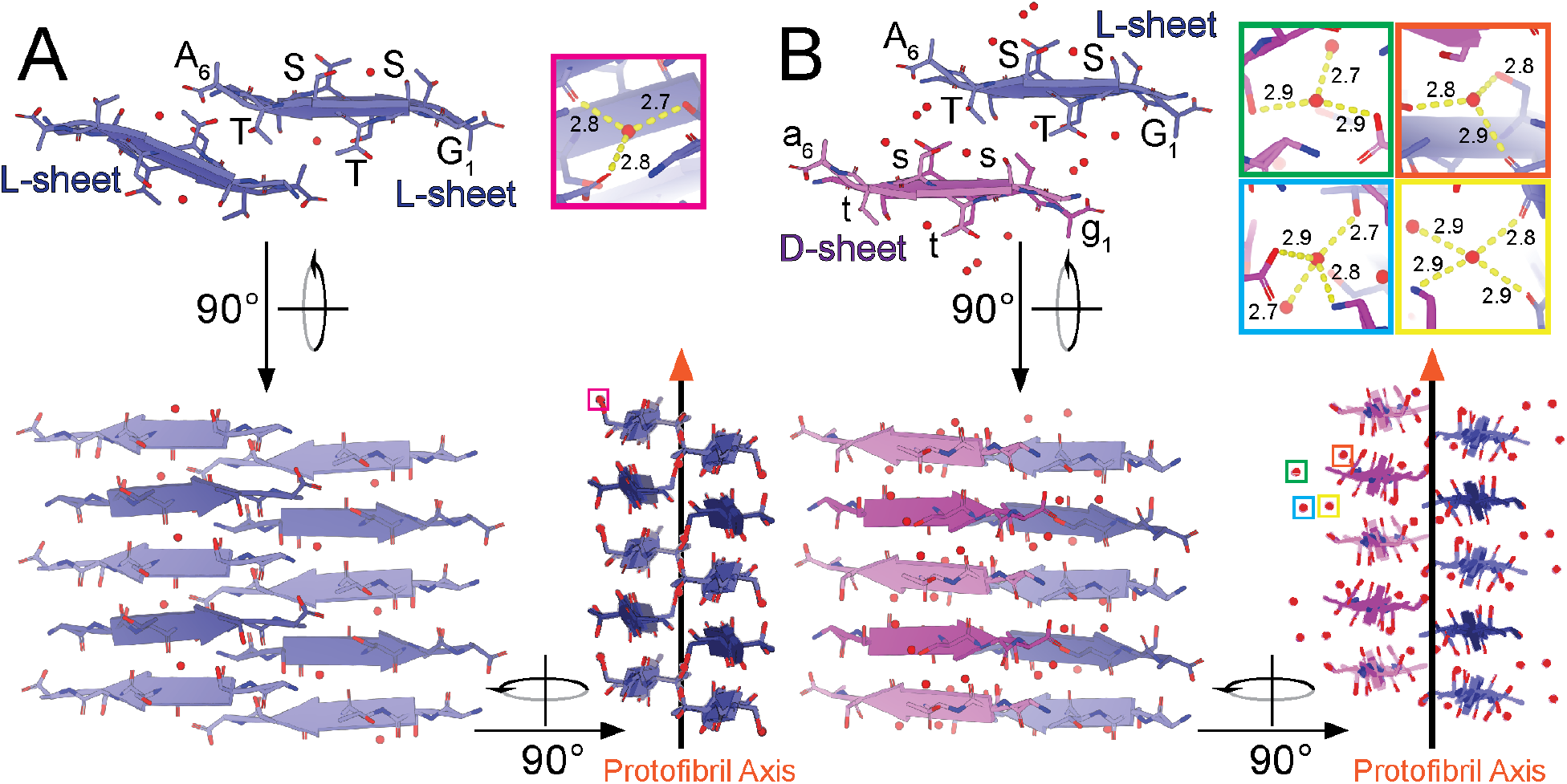
Views of protofibrils of L-GSTSTA (A) and racemic GSTSTA (B) represented by pair of sheets with a view down the protofibril axis; both structures derived by x-ray diffraction. A 90° rotation shows a side view of the protofibril with strands stacked along each sheet in an antiparallel fashion. Another 90° rotation shows a side view of the protofibril along the strand axis, showing a buckling of each sheet due to the tilting of strands away from or toward the protofibril axis. Chains colored such that blue represent L-peptides while magenta represents D-peptides. Lighter and darker shades of each color differentiate the orientation of strands within a sheet. Ordered waters found in each asymmetric unit are indicated by colored squares that correspond to insets of matching colors. Insets show magnified views of each water molecule with hydrogen bonds represented by the yellow dashed lines, labelled with their corresponding distances in Å

**Figure S14.**
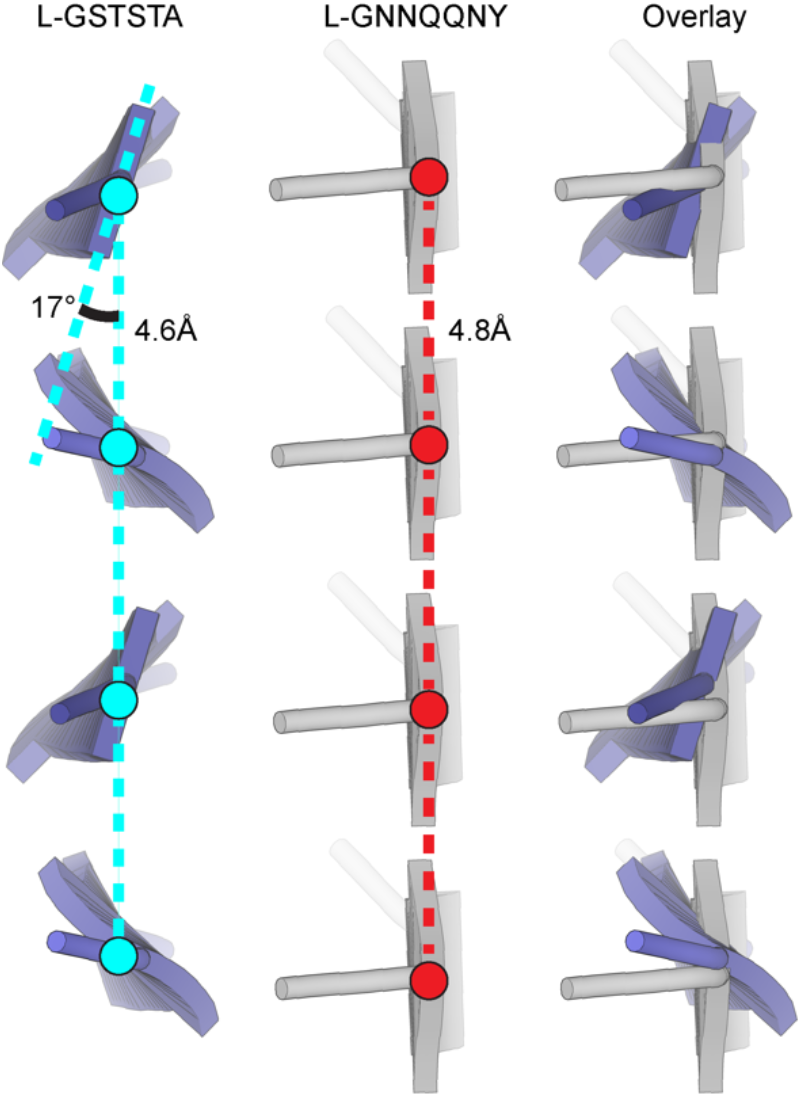
Beta sheets in the structure of L-GSTSTA have an inter-strand distance of 4.6Å with amide carbonyls angled away from the protofibril axis by approximately 17° (left). Canonical beta sheets formed by the yeast prion segment L-GNNQNNY (RCSB PDB: 1YJP) show inter-strand distance of 4.8Å and a near 0° deviation of amide hydrogen bonding down the protofibril axis (center) (Nelson *et al*., 2005). An overlay (right) illustrates compaction of the L-GSTSTA sheet along its length compared to a sheet formed by L-GNNQQNY.

**Figure S15.**
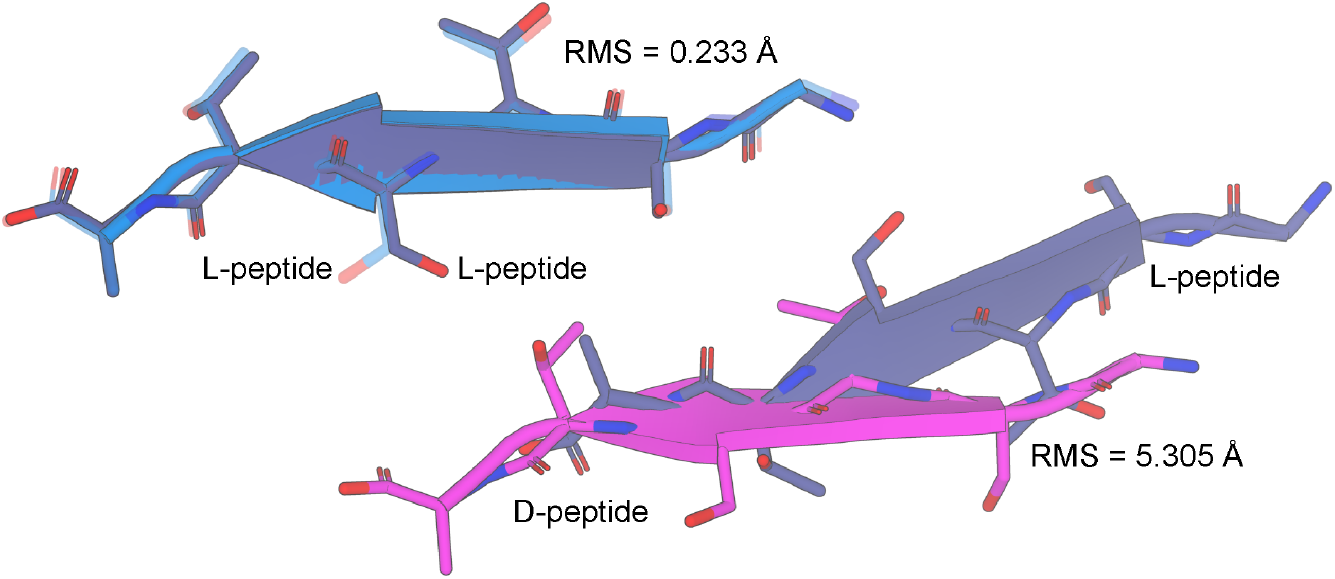
Pairs of mated strands representing the homochiral and racemic protofibrils of GSTSTA. Alignment of these protofibrils based on a common L-GSTSTA sheet shows a displacement of their paired sheet. The RMSD between the common L-GSTSTA sheets is 0.23 Å, while that between the pairing L and D sheets of the homochiral and racemic protofibrils is 5.3 Å. L-GSTSTA strands are colored blue and purple while D-GSTSTA is colored magenta.

**Figure S16.**
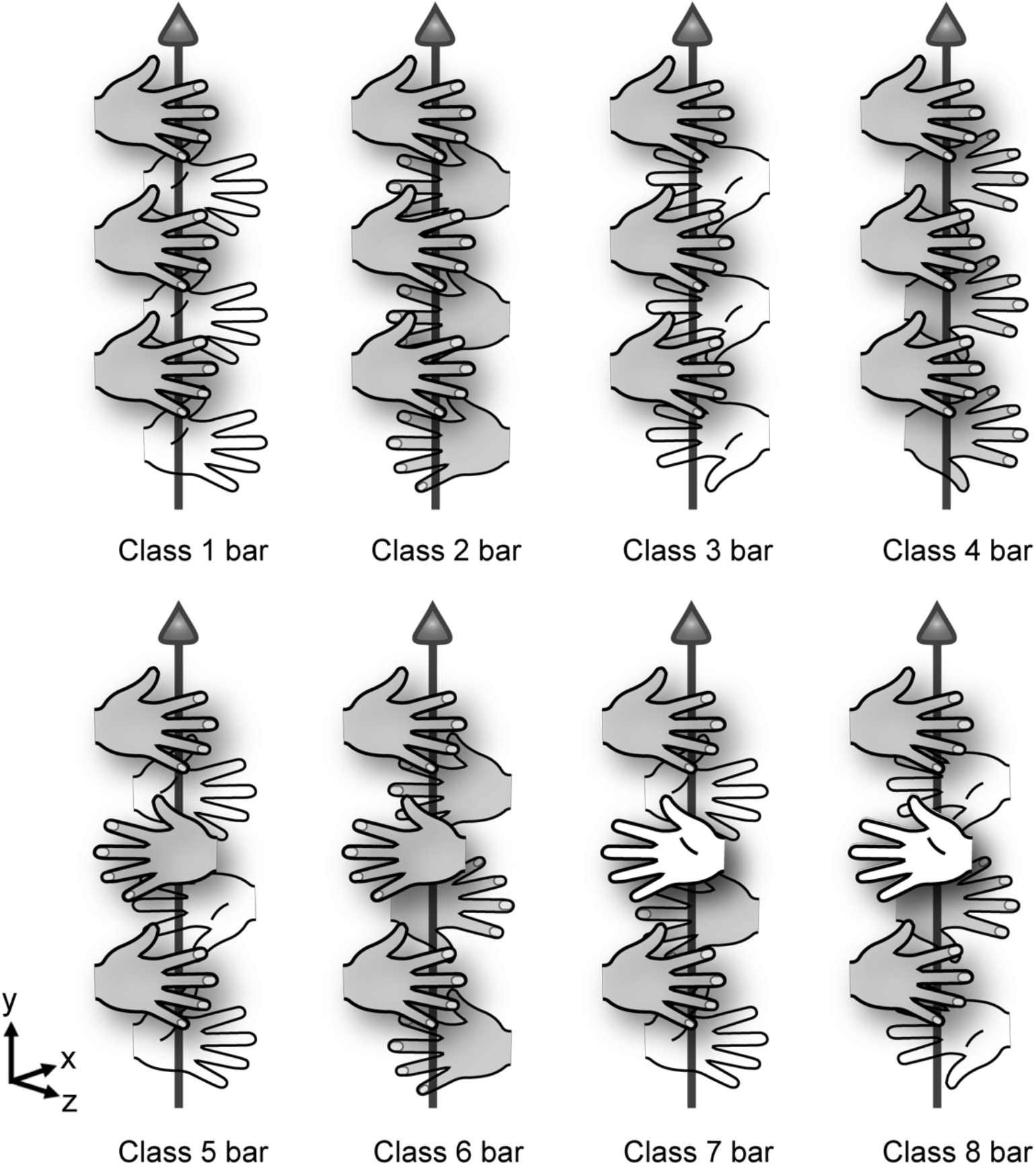
Eight new potential steric zipper classes are enabled by racemic assemblies. All new classes are based on those originally described in Sawaya *et al*. (Sawaya *et al*., 2007) but now contain a mirror plane, a glide plane, or an inversion center. Strands are represented by left and right hands, each equivalent to the enantiomers present in a zipper class. The asymmetry of side chains on either side of a strand is portrayed by the palm and back of each hand. The up or down orientation of the thumbs and the direction in which the fingers point indicates the direction of each strand. An arrow indicates the axis of fibril growth, which here is coincident with the y direction. While additional symmetry classes with racemic mixtures are possible, only eight are illustrated here.

